# Cilium proteomics reveals Numb as a positive regulator of the Hedgehog signaling pathway

**DOI:** 10.1101/2022.10.10.511655

**Authors:** Xiaoliang Liu, Patricia T. Yam, Sabrina Schlienger, Eva Cai, Jingyi Zhang, Wei-Ju Chen, Oscar Torres Gutierrez, Vanesa Jimenez Amilburu, Vasanth Ramamurthy, Alice Y. Ting, Tess C. Branon, Michel Cayouette, Risako Gen, Tessa Marks, Jennifer H. Kong, Frédéric Charron, Xuecai Ge

## Abstract

The transduction of Hedgehog (Hh) signaling relies on the primary cilium, a cell surface organelle serving as a signaling hub for the cell. Using proximity labeling and quantitative proteomics, we identified Numb as a new ciliary protein that positively regulates Hh signaling. Numb localizes to the ciliary pocket and acts as an endocytic adaptor to incorporate Ptch1 into clathrin-coated vesicles, thereby promoting Ptch1 exit from the cilium, a key step in Hh signaling activation. Numb loss hampers Sonic Hedgehog (Shh)-induced Ptch1 departure from the cilium, resulting in reduced activation of Hh signaling. Numb loss in spinal neural progenitors reduces Shh-induced differentiation into Nkx2.2-positive progenitors, a process reliant on high Hh signaling activity. Genetic ablation of Numb in the developing cerebellum impaired the proliferation of granule cell precursors, a Hh-dependent process, resulting in reduced cerebellar size. This study highlights Numb as a critical regulator of Ptch1 levels in the cilium during Hh signal activation and demonstrates the key role of ciliary pocket-mediated endocytosis in modulating the transduction of cell signaling.

## INTRODUCTION

The primary cilium serves as a cellular antenna that senses molecules and conveys their signal from the extracellular milieu into the cell^1, 2^. Hedgehog (Hh) signaling was the first signaling pathway discovered to rely on the primary cilium^3^. Hh signal transduction is carried out by the dynamic transport of signal transducers into and out of the cilium. In the basal state, the Hh receptor Ptch1 resides in the cilium where it prevents the accumulation and activation of Smo in the cilium^4^. Upon Shh stimulation, Ptch1 exits the cilium. Reciprocally, Smo accumulates and is activated in the cilium. Active Smo triggers a signaling cascade that eventually activates Gli transcription factors which then turn on the transcription of Hh target genes^5–7^. During this process, Ptch1 exit from the cilium acts as a key step to initiate Hh signaling.

Recent studies suggest that Ptch1 acts as a sterol transporter to control local sterol components, such as cholesterols, in the cilium^8–12^. Binding to certain oxysterols is required for Smo activation^13, 14^. Therefore, Ptch1 is believed to inhibit Smo by restricting Smo’s access to activating sterols in the ciliary membrane. The clearance of Ptch1 from the cilium correlates with Shh-mediated pathway activation and is necessary for maximal activation of Hh signaling^4, 15^. However, the endocytic machinery that mediates Ptch1 exit from the cilium remains largely unknown. Yue et al. reported that Ptch1, a substrate of the E3 ubiquitin ligases Smurf1 and Smurf2, colocalizes extensively at the cytoplasmic membrane with the two E3 ligases^16^. Ubiquitination of Ptch1 promotes its lysosomal degradation following caveola-mediated endocytosis. However, this mechanism depicts Ptch1 endocytosis from the cytoplasmic membrane, and may not apply to the process of Ptch1 exiting from the cilium^17^. Another recent study showed that Ptch1 in commissural neuron growth cones is internalized via Numb, a well-established adaptor for clathrin-mediated endocytosis. This process is required for non-canonical Hh signaling and growth cone turning up Shh gradients^18^. Whether Numb acts as an endocytic adaptor to mediate Ptch1 departure from the cilium in canonical Hh signaling remains to be determined.

A critical approach to understanding signaling mechanisms in the cilium is to catalog the dynamic protein components in the cilium over the course of signal transduction. This has been technically challenging due to the miniature size of the cilium, which accounts for about 1/10,000 of the cytoplasmic volume. Proximity-based labeling tools provide a powerful method to specifically label and identify ciliary proteins. APEX and its variant APEX2, engineered peroxidases, have been used to specifically label proteins in the cilium^19–21^. These enzymes were targeted to the primary cilia via distinct ciliary proteins. Intriguingly, when the enzyme was targeted to different sub-ciliary domains by fusing to distinct “tagging” proteins, unique cohorts of cilium proteins were discovered.

In this study, we leveraged a mechanistically different proximity labeling tool, TurboID, to plot ciliary proteins over the course of Hh signal transduction. TurboID is an engineered promiscuous biotin ligase that labels neighboring proteins and avoids the need for toxic labelling conditions dependent on hydrogen peroxide^22^. Using a truncated form of the membrane-associated ciliary protein Arl13b to tag TurboID, our ciliary proteome recovered a wide array of new ciliary protein candidates. Among the new ciliary candidates, we investigated the mechanistic roles of Numb, a protein previously shown to involve in signaling regulation and cell fate specification^23–26^. Interestingly, we found that Numb localizes to the ciliary pocket, a cytoplasmic membrane region at the base of the cilium that “folds back” to ensheath the cilium. Moreover, we found that Numb interacts with Ptch1 and recruits Ptch1 to clathrin coated vesicles (CCVs), thereby promoting Ptch1 exit from the cilium. Numb loss in NIH3T3 cells reduces the plateau levels of Hh signaling elicited by high concentrations of Shh. Further, in mouse embryonic stem cell (ESC)-derived spinal cord neural progenitors (NPCs), Numb depletion blocks cell differentiation into an identity that requires high Hh activity. Genetic ablation of Numb in the developing cerebellum decreased the proliferation of GCPs, a Hh-signaling dependent process, resulting in reduced cerebellar size. Overall, our study highlights a new profile of membrane-associated proteins in the cilium, and demonstrates the molecular role of Numb as a critical regulator of endocytosis in the ciliary pocket, a process crucial for the transduction of cilium-mediated cell signaling.

## RESULTS

### Cilium-TurboID selectively biotinylates ciliary proteins

Many signaling proteins are membrane proteins or recruited to the membrane during signal transduction. To label these proteins in the primary cilium, we directed TurboID to the juxtamembrane region of the cilium. We chose Arl13b to fuse with TurboID since Arl13b is anchored to the ciliary membrane via lipid modification^27, 28^, maximizing the chance of protein biotinylation in the juxtamembrane region. As reported before, overexpressing full-length Arl13b leads to markedly elongated cilia in NIH3T3 cells^29^ (Supplementary Fig. 1a). To determine which domains of Arl13b could retain ciliary localization without altering cilium length, we generated a series of truncated Arl13b constructs and transiently expressed them in NIH3T3 cells (Fig. 1a and Supplementary Fig. 1a). We found that the N+RVEP+PR construct fits this requirement. This construct contains the N-terminal amphipathic helix (containing a palmitoylation modification for membrane association), the previously reported sequence RVEP that is critical for cilium targeting, and the Proline Rich domain at the C-terminus^30^ (Fig. 1a). In contrast, the construct containing only the PR domain showed no ciliary localization (Supplementary Fig. 1a-b).

**Figure 1.**
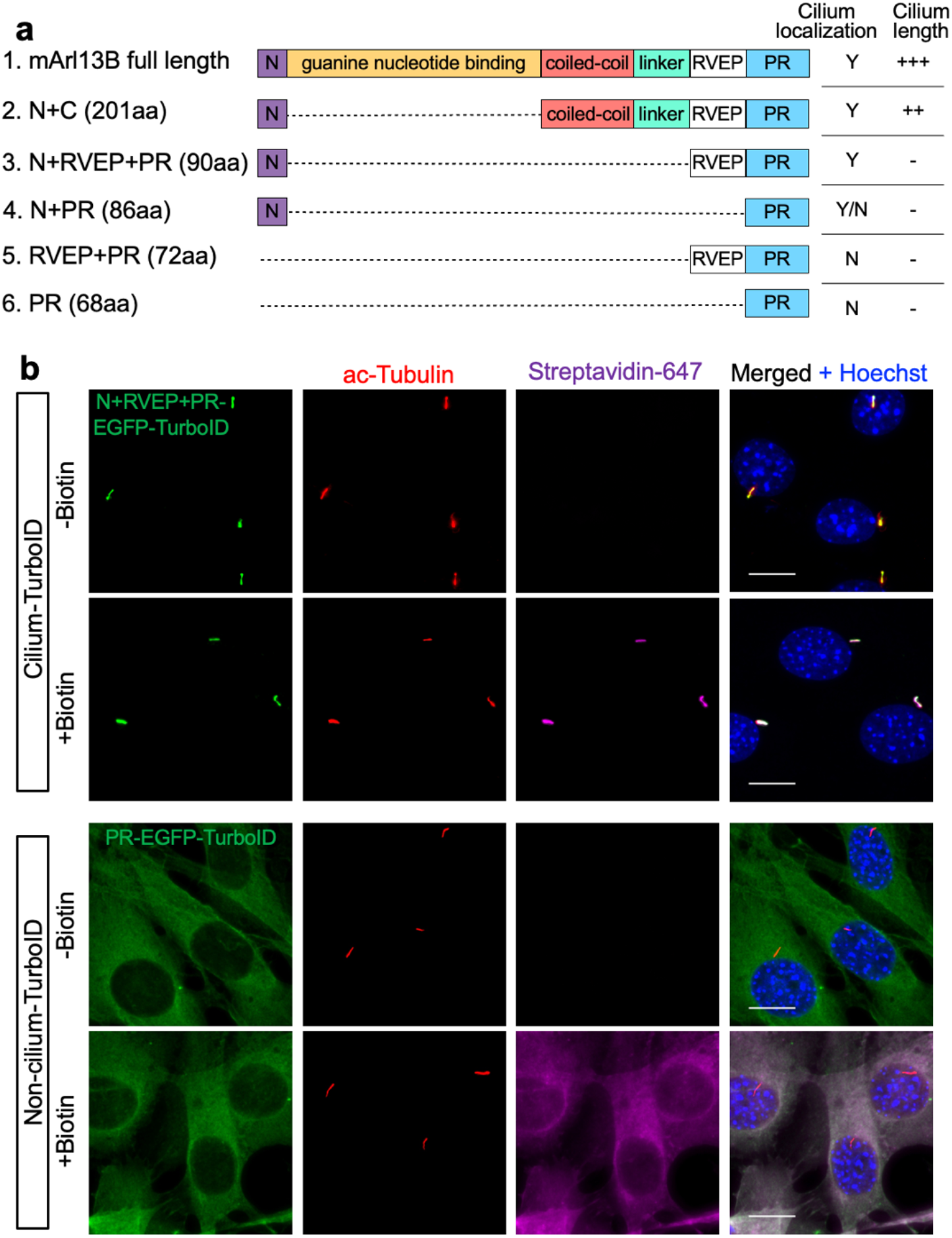
Cilium-TurboID selectively biotinylates ciliary proteins. (**a**) Schematics of the truncated Arl13b constructs used. Y and N indicate whether the corresponding construct localizes to the cilium or not, respectively. Y/N: this construct localizes to both the cilium and the cytosol. “+”: the degree of cilium elongation in NIH3T3 cells transiently transfected with the corresponding constructs. “-”: cilium length is not impacted. (**b**) Immunofluorescence of stable NIH3T3 cell lines expressing cilium-TurboID (N+RVEP-PR-EGFP-TurboID) or non-cilium-TurboID (PR-EGFP-TurboID) before and after biotin labeling. Cells were serum-starved for 24 h, incubated with 50 μΜ biotin for 10 min at 37°C, and fixed for immunostaining. The cilium is labeled by immunostaining for acetylated-tubulin (red, ac-tubulin), and the biotinylated proteins were detected via streptavidin-Alexa Fluor 647 (magenta). Cilium-TurboID and non-cilium TurboID were detected by immunostaining for EGFP (green). Scale bars, 10 µm in all panels.

We next fused TurboID to N+RVEP+PR (cilium-TurboID), and to the PR domain (non-cilium-TurboID) as a control for cytosolic proteins. We then generated stable NIH3T3 cell lines that express cilium- or non-cilium-TurboID via an integral lentiviral system. Confocal microscopy confirmed that cilium-TurboID is exclusively localized to the primary cilium, whereas non-cilium-TurboID is diffusely distributed in the cytoplasm (Fig. 1b). Furthermore, 10 min labeling with biotin selectively labeled proteins in the cilium in cilium-TurboID cells, whereas cytosolic proteins were labeled in non-cilium-TurboID cells (Fig. 1b).

To ensure that the transgenes did not impair cilium function, we measured cilium length and Hh signaling activity in the stable cell lines. We found that the cilium lengths are comparable between WT (parental NIH3T3 cells), cilium-TurboID and non-cilium-TurboID cells (Supplementary Fig. 2a). We then determined the Hh signaling activity after the cells were stimulated with Shh. The Hh signaling activity was evaluated by the transcript levels of *Gli1*, a Hh target gene. We found that Shh-induced *Gli1* levels were comparable between WT and stable cell lines (Supplementary Fig. 2b). Further, Shh-induced cilium transport of Smo and Gli2 are indistinguishable between WT and the stable cell lines (Supplementary Fig. 2c-f). In summary, the cilium morphology and Hh signaling are normal in these stable cell lines. Thus, we developed a method to exclusively target TurboID to the juxtamembrane microdomain in the primary cilium without impacting cilium functions.

### Quantitative proteomics with cilium-TurboID highlighted membrane-associated ciliary proteins

We then employed the stable cell lines to label and isolate endogenous ciliary proteins. We labeled cells with biotin for 10 min and captured the biotinylated proteins with streptavidin beads^22^. Proteins were analyzed by Western blotting (Fig. 2a). Streptavidin-HRP detection of the biotinylated proteins showed that biotin labeling drastically increased the overall levels of biotinylated proteins captured by the streptavidin beads. As a positive control, we examined PDGFRα, a receptor known to be localized to the cilium. As expected, PDGFRα was pulled down from cilium-TurboID cells, but not non-cilium-TurboID cells (Fig. 2a). These results confirm that cilium-TurboID is effective in labeling proteins in the cilium.

**Figure 2.**
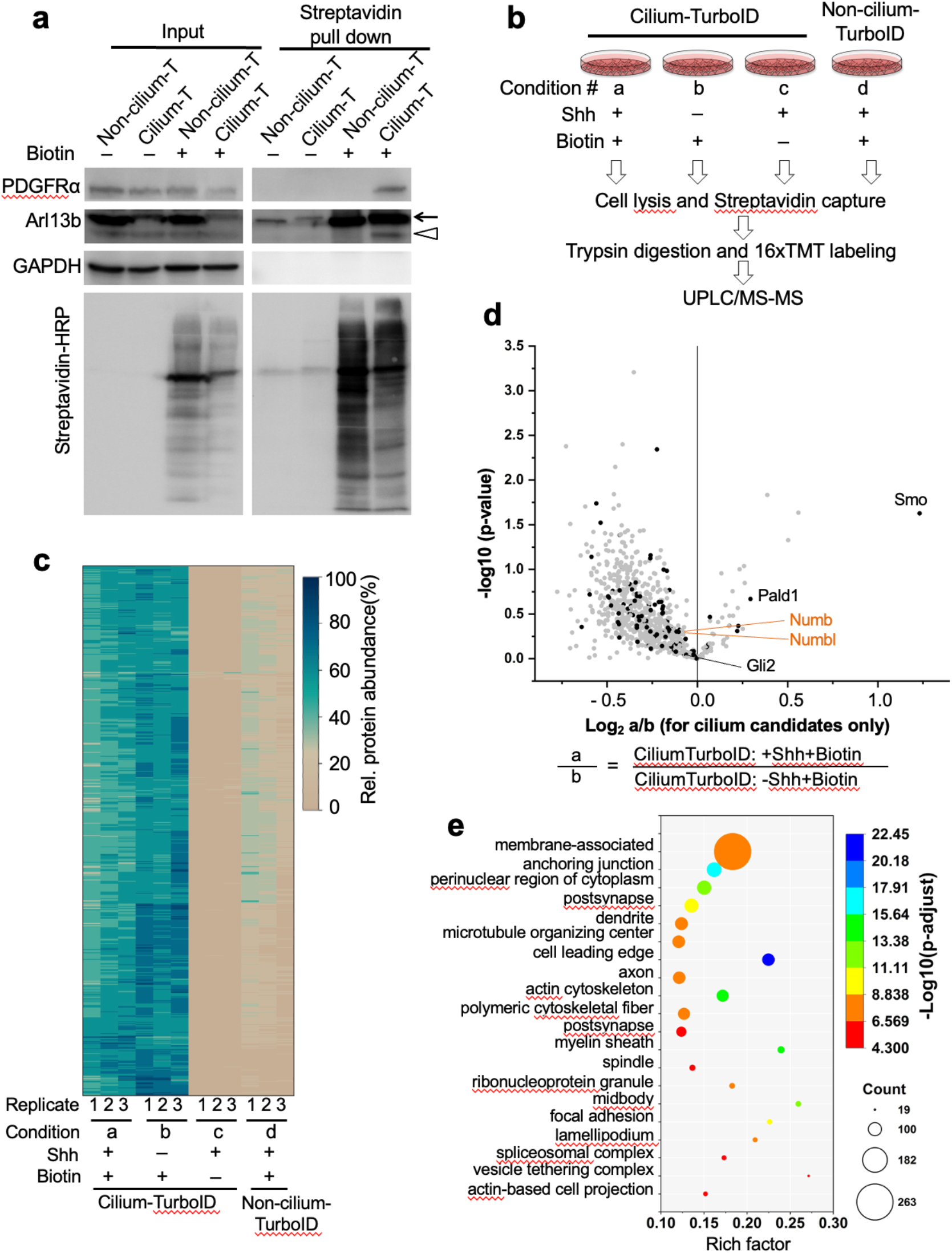
Quantitative proteomics with cilium-TurboID identified novel ciliary proteins. (**a**) Biotinylated proteins from cell lysates of cilium-TurboID and non-cilium-TurboID cells were isolated by streptavidin beads and analyzed by Western blotting. The overall biotinylated proteins were detected by streptavidin-HRP. PDGFRα is a receptor known to localize to the cilium. In Arl13b blot, the arrow points to the transgene, and the triangle points to endogenous Arl13b. (**b**) Design and workflow of the cilium proteomics experiment. Samples were prepared in triplicate. After TMT labeling, all samples were pooled together before being fractionated by UPLC and analyzed by mass spectrometry. (**c**) Heatmap of the ciliary abundance of the 800 ciliary candidates. Clustering of the relative abundances of each identified protein (rows) in individual samples (columns) was performed based on Ward’s minimum variance method. The relative abundance was calculated within one set of samples that contains all four conditions. For example, a1, b1, c1, d1 belong to one set, and the relative abundance is calculated as a1/(a1+b1+c1+d1) etc. Right, the color scheme of relative abundances (percentage). (**d**) Volcano plot of statistical significance versus the average ratio of protein enrichment for all ciliary candidates. The ratio was calculated as TMT signal intensity of Shh-treated samples (a) compared to none-Shh samples (b) in the presence of biotin labeling. The analytical procedure for identifying ciliary candidates from the mass spectrometry results is described in Fig. S3A-D. The previously reported ciliary proteins are highlighted as black dots. (**e**) GO enrichment analysis of cellular components was plotted according to the rich factor in Metascape. The top 20 enriched cellular components are represented in the scatter plot.

Next, we designed a quantitative proteomic experiment to capture the dynamic protein transport in the cilium in response to Shh stimulation. To control for background biotinylation, we included conditions that omitted either Shh stimulation or biotin labeling. To control for non-ciliary protein biotinylation by TurboID, we collected samples from stimulated and labeled non-cilium-TurboID cells (Fig. 2b). We prepared each condition with three independent replicates. After on-beads digestion, each sample was labeled with one channel of the 16plex TMT labeling kit. A reference channel was included in which equal volumes from each of the 12 samples were pooled. After that, the 13 samples were multiplexed and analyzed by liquid chromatography (UPLC)/MS-MS (Fig. 2b; channel distribution in supplementary figure 3a).

For data analysis, we defined the relative protein abundance as the ratio of normalized abundance in each channel over the reference channel. Only proteins with more than 7 unique peptides were used in data analysis. We determined candidate ciliary proteins via statistical analyses of the relative enrichment between the cilium-TurboID and the non-cilium-TurboID datasets. To be scored as ciliary proteins in the absence of Shh stimulation, candidates had to fulfill four criteria: 1) greater than 2-fold enrichment (TMT ratio > 2) in the labeled cilium-TurboID samples over the non-labeled cilium-TurboID samples, 2) and over the labeled non-cilium-TurboID samples; 3) statistically significant (p value < 0.05) enrichment in the labeled cilium-TurboID samples versus the non-labeled cilium-TurboID samples, and 4) versus the labeled non-cilium-TurboID samples (Supplementary Fig. 3b, c). 788 proteins meet these criteria (Table S1). To be scored as ciliary proteins after Shh stimulation, candidates had to fulfill the same four criteria, comparing between the labeled Shh-stimulated cilium-TurboID samples and the non-labeled Shh-stimulated cilium-TurboID samples, and the labeled Shh-stimulated non-cilium-TurboID samples (Supplementary Fig. 3d, e). 574 proteins meet these criteria (Table S1). Finally, we took the union of the above two categories and defined these proteins as ciliary candidates regardless of Shh stimulation. We obtained 800 such ciliary candidates in total (Table S2). Correlation of biological replicates and a heatmap of each protein’s relative abundance in the 12 samples demonstrated a high reproducibility across the triplicates (Fig. 2c; Supplementary Fig. 4a-d).

We then compared the intensity of the above identified ciliary candidates between the Shh-stimulated and unstimulated conditions (Fig. 2d, Table S2). We identified 30 proteins that show statistically significant changes (TMT ratio > 1.2 or < 0.83, p < 0.05). Among them is the Hh transducer Smo. It is important to note that a lack of statistical significance does not imply that the candidates do not respond to Hh signaling, owing to a variety of reasons such as 1) the low abundance of the protein in the cilium, and 2) preferential biotinylation of proteins in juxtamembrane subdomain in our experimental settings. Indeed, a few proteins known to translocate to the cilium in response to Hh activation show larger p value, such as Gli2 and PALD1^20^.

Among the total ciliary candidates, 108 have been reported before to localize to the cilium (Supplementary Fig. 4e). Analysis of the ciliary candidates with enrichment of Gene Ontology (GO) terms in Metascape shows that the largest cohort are membrane-associated proteins (Fig. 2e). These results validated our strategy of targeting TurboID to the ciliary juxtamembrane domain.

### Numb localizes to the ciliary pocket and clathrin-coated vesicles (CCVs)

Among the newly identified ciliary candidates, Numb is particularly interesting. *Drosophila* Numb has important roles in asymmetric cell division^23^, and mammalian Numb regulates neurogenesis in the developing brain^24–26^. However, the cilium localization of Numb has never been described before. Our proteomic results revealed Numb as a potential ciliary protein. The mass spectrometry intensities of Numb in the cilium display a reduced tendency following Shh stimulation, although no statistically significant changes are observed (Fig. 2d). To validate its ciliary localization, we expressed V5 tagged Numb in NIH3T3 cells. We found that in addition to its reported distribution in the cytosol as puncta^31, 32^, Numb-V5 also overlaps with the ciliary marker Arl13b in 33% of transfected cells (Fig. 3a). However, the Numb immunofluorescence signal is not distributed along the entire cilium; rather, it is restricted to the lower section of the cilium.

**Figure 3.**
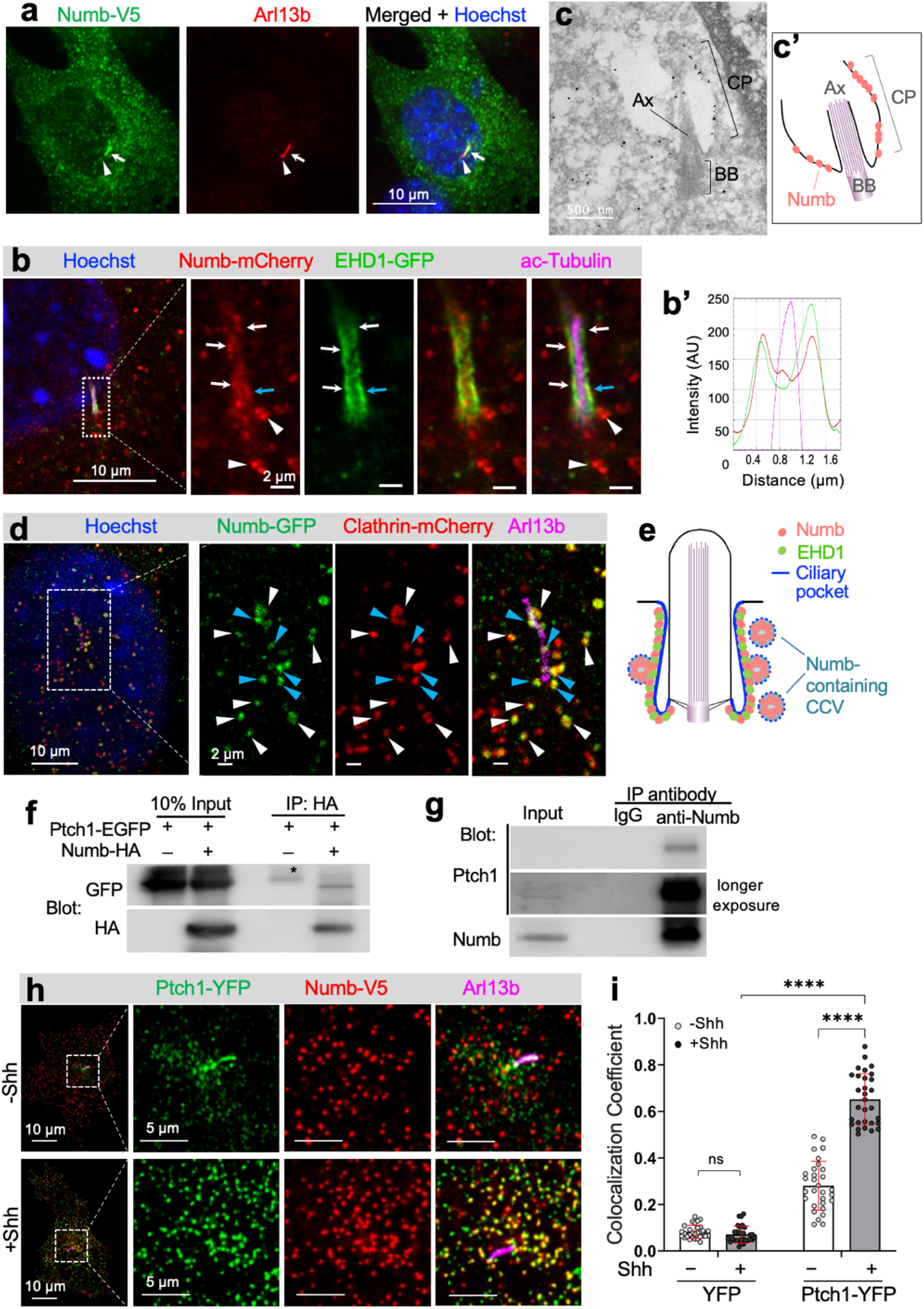
Numb localizes to the ciliary pocket and incorporates Ptch1 into clathrin coated vesicles. (**a**) Numb localizes to the lower section of the primary cilium when expressed in NIH3T3 cells. Cilium is highlighted by Arl13b staining. White arrow points to the lower section of the cilium; white triangle points to the ciliary tip where Numb immunofluorescence are absent. (**b**) Numb co-localizes with EHD1 at the ciliary pocket. NIH3T3 cells were co-transfected with Numb-mCherry and EHD1-GFP, subjected to expansion, and imaged with Airyscan microscopy. The displayed image corresponds to a single focal plane, encompassing only the lower segment of the cilium axoneme. The cilium axoneme is labeled with acetylated-tubulin (Magenta). Arrows point to the ciliary pocket where Numb and EHD1 exhibit colocalization; white triangles indicate Numb-containing puncta located adjacent to the cilium. (**b’**) The linescan was performed at the position marked by the cyan arrow to illustrate the colocalization of Numb with EHD1 in the ciliary pocket. Red: Numb-mCherry, green: EHD1-GFP, magenta: ac-tubulin. (**c**) Numb exhibits localization to the ciliary pocket in ImmunoEM. NIH3T3 cells expressing Numb-HA were stained with anti-HA antibody, followed by the secondary antibody conjugated to 1.4 nm nanogold. The nanogold signal was enhanced by GoldEnhance kit. Ciliary pocket (CP). Ax, axoneme; BB, basal body. Scale bar, 500 nm. (**d**) Numb localizes to clathrin coated vesicles (CCVs). NIH3T3 cells were co-transfected with Numb-GFP and clathrin-mCherry, subjected to expansion, and imaged with Airyscan microscopy. The displayed image corresponds to a single focal plane, highlighting CCVs near the cilium base. The cilium membrane is labeled by Arl13b (magenta). White triangles point to the Numb-containing CCVs. Cyan triangles point to Numb-containing CCVs still in conjunction with the ciliary pocket, suggesting that they are emerging CCVs from the ciliary pocket. (**e**) The diagram illustrates Numb’s localization to the ciliary pocket and to CCVs situated in close proximity to the ciliary pocket. (**f**) Ptch1-EGFP and Numb-HA were expressed in 293T cells. Numb-HA was immunoprecipitated with an anti-HA antibody, and Ptch1-EGFP was co-immunoprecipitated with Numb-HA. * indicates a non-specific band. (**g**) In NIH3T3 cells treated with Shh, endogenous Ptch1 was co-immunoprecipitated with Numb via a Numb antibody. (**h**) Ptch1 is incorporated into Numb-containing CCVs near the ciliary pocket. NIH3T3 cells were co-transfected with Numb-V5 and either Ptch1-YFP or YFP (images for YFP in supplementary figure 7). Cells were then treated with Shh or vehicle for 30 min and fixed for immunostaining. The cilium is labeled by Arl13b (magenta). Enlarge areas are regions surrounding the cilium. (**i**) Colocalization Coefficient of Numb-V5 and Ptch1-YFP or YFP. Manders’ Colocalization Coefficients (MCC) analysis was performed using BIOP JACoP plug-in in ImageJ. A total of 30 cells were quantified for each experimental condition. Statistics: Two-way ANOVA with multiple comparisons (Tukey test), ****, p < 0.0001; ns, not significant. All error bars represent standard deviation (SD).

The localization pattern of Numb in the cilium does not resemble any of the known cilium subdomains, such as the transition zone^33^, inversin zone^34^, or Evc zone^35^. Rather, it resembles the recently characterized ciliary pocket, a cytoplasmic membrane region at the bottom of the cilium that often folds back to ensheath the bottom of the primary cilium^36, 37^. To characterize Numb’s localization in the ciliary pocket, we co-transfected NIH3T3 cells with Numb-mCherry and EHD1-GFP, a protein known to localize to the ciliary pocket^38^(Fig. 3b). We then performed expansion microscopy, a procedure that physically expands the samples in an isotropic fashion to allow nanoscale resolution^39, 40^. After expansion, cells were imaged with Airyscan microscopy. The cilium axoneme is highlighted with acetylated-tubulin staining. We found that Numb overlaps with EHD1 at the lower part of the cilium in the region flanking the cilium axoneme (Fig. 3b, b’), suggesting that Numb localizes to the ciliary pocket. To further confirm this result, we performed immuno-electron microscopy in NIH3T3 cells expressing HA-tagged Numb. Notably, extensively electron-dense staining is observed in the ciliary pocket membrane (Fig. 3c-c’). Together, our results indicate that Numb localizes to the ciliary pocket.

In addition to the ciliary pocket, Numb immunofluorescence appeared as discrete puncta in the cytosol (white triangles in Fig 3b). Since Numb is reported to be involved in clathrin-mediated endocytosis^32^, these puncta are likely clathrin-coated vesicles (CCVs). To test this, we co-transfected NIH3T3 cells with Numb-GFP and clathrin-mCherry, and imaged cells via expansion microscopy. At the focal plane focusing on the bottom of the cilium, we found that at the region adjacent to the cilium, all Numb-containing puncta are clathrin positive, suggesting that these puncta are CCVs (Fig. 3d). Further, multiple double positive puncta are still in close proximity to the cilium (Fig. 3d, cyan triangles), indicating that these puncta are very likely CCVs emerging from the ciliary pocket^37^. Taken together, Numb localizes to the ciliary pocket and to CCVs derived from the ciliary pocket (Fig. 3e).

### Numb interacts with Ptch1 and incorporates Ptch1 into CCVs

The ciliary pocket is believed to play a crucial role in the endocytic control of ciliary proteins^36, 37, 41^. Our findings indicate that Numb is not only present in the ciliary pocket but also associated with CCVs linked to the ciliary pocket. Given Numb’s established function as an adaptor that connects endocytosed cargoes with adaptins in the clathrin-mediated endocytic machinery^42, 43^, it is plausible that Numb may participate in endocytosis at the ciliary pocket. During Hh signal transduction, Ptch1 exit from the cilium is a critical step to ensure the maximal activation of Hh signaling. We hypothesized that Numb recruits Ptch1 to CCVs at the ciliary pocket, thereby mediating Ptch1 exit from the cilium.

To test our hypothesis, we first examined whether Ptch1 and Numb interact with each other in cells. In 293T cells, Ptch1-GFP co-immunoprecipitated with Numb-HA (Fig. 3f). We then tested whether endogenous Ptch1 and Numb form complexes with each other in ciliated cells. We immunoprecipitated endogenous Numb from lysates of NIH3T3 cells and found that endogenous Ptch1 co-immunoprecipitated with Numb (Fig. 3g). Therefore, Numb and Ptch1 form complexes with each other. To determine which domain mediates this interaction, we employed Numb truncated proteins in co-immunoprecipitation assays. We found that deletion of either the N-terminus (aa1-25) or the PTB domain (aa26-173) abolishes the interaction with Ptch1. Meanwhile, the N+PTB domain (aa1-173) is sufficient for interaction with Ptch1 (Supplementary Fig. 5a, b). These results indicate that Numb interacts with Ptch1 via its N-terminal and PTB domain.

Next, we determined whether Ptch1 is incorporated into Numb-containing CCVs. We co-expressed Numb and Ptch1 in NIH3T3 cells via lentiviruses that allow low expression levels in order to recapitulate the behavior of endogenous proteins. In the absence of Shh, only a limited number of Numb-positive puncta contain Ptch1-YFP. In contrast, upon Shh stimulation, there is a notable increase in the colocalization of Numb and Ptch1 within the puncta at the surrounding region of the cilium. The colocalization coefficient rises from 0.28 ± 0.1 (without Shh) to 0.65 ± 0.1 (with Shh) (Fig. 3h-i). As a control, we co-expressed YFP with Numb-V5, and did not observe any significant overlap between Numb and YFP (Supplementary Fig. 6a). These results suggest that Shh stimulation promotes Ptch1 incorporation into Numb-containing CCVs.

Taken together, our results suggest that Numb interacts with Ptch1 and incorporates Ptch1 into CCVs to promote Ptch1 internalization from the ciliary pocket upon Shh stimulation.

### Numb is required for Shh-induced Ptch1 exit from the cilium

To determine Numb’s role in Ptch1 exit from the cilium, we generated Numb knockout cells via CRISPR/Cas9 in NIH3T3 cells. We used two guide RNAs to target the first and third exons of Numb, and isolated 14 single cell colonies of Numb knockout (KO) cells (Supplementary Fig. 7a). We then validated biallelic indel mutations that eliminate Numb expression in two of the cell clones (Supplementary Fig. 7b, c). In these two clones, Numb protein was undetectable in Western blotting (Fig. 4a). We used these two cell clones in the subsequent experiments.

**Figure 4.**
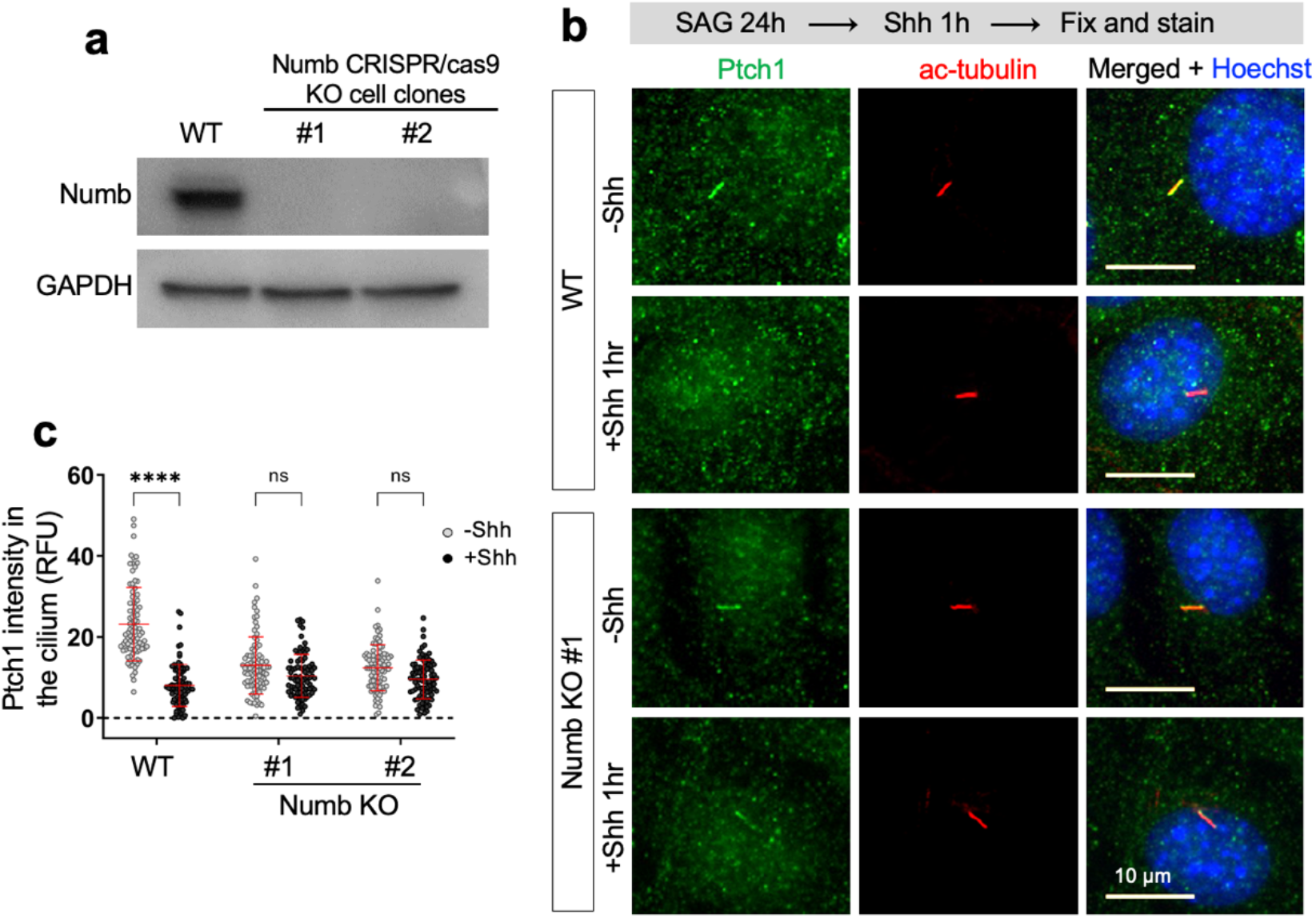
Numb is required for Shh-induced Ptch1 exit from the cilium. (**a**) NIH3T3 cells underwent Numb knockout (KO) using CRISPR/Cas9 technique. The resulting cell clones were isolated individually. Western blot analysis confirmed the absence of Numb protein in the Numb CRISPR/Cas9 knockout cell clones. (**b**) Wild type (WT) or Numb KO cells were pre-treated with SAG for 24 h to induce Ptch1 expression and cilium accumulation, followed by a 1h’s Shh stimulation, and subsequently fixed for immunofluorescence staining. The cilia were marked by ac-tubulin (red). (**c**) Quantification of Ptch1 intensity in the cilium. Data are shown as mean ± SD. A total of 50 cells were quantified for each experimental condition. Statistics: Two-way ANOVA with multiple comparisons (Tukey test). ****, p < 0.0001. ns, not significant.

We next determined the impact of Numb loss on the exit of Ptch1 from the cilium. As described before^4^, in unstimulated cells, Ptch1 in the cilium is below detection threshold of immunostaining, but overnight SAG treatment induces Ptch1 expression and localization to the cilium (Fig. 4b). Following that, Shh stimulation triggers Ptch1 exit from the cilium. In WT cells, Ptch1 intensity significantly decreased in the cilium after Shh stimulation. In contrast, in Numb KO cells, the ciliary Ptch1 intensity remained comparable before and after Shh stimulation (Fig. 4b, c). In a parallel experiment, we expressed low levels of Ptch1-YFP via lentivirus in WT or Numb KO cells. Shh effectively induced Ptch1-YFP exit from the cilia in WT cells. In contrast, Ptch1-YFP levels remained unchanged before and after Shh in Numb KO cells (Supplementary Fig. 8a, b). To exclude the potential impact of Numb loss on the formation of the ciliary pocket, we visualized the pocket after expansion microscopy. The morphology of the ciliary pocket, marked by EHD1-GFP, showed no discernible differences between the WT and Numb KO cells (Supplementary Fig. 8c). In conclusion, Numb loss hinders Ptch1 exiting from the cilium.

### Numb loss diminishes the plateau levels of Hh signaling

In the absence of Hh stimulation, Ptch1 in the cilium suppresses Smo activation (Fig. 5a). Hh stimulation induces Ptch1 exit from the cilium, which is critical for the activation of Hh signaling. We predicted that without Numb, Hh signaling would be suppressed due to impaired Ptch1 exit from the cilium. To test this, we evaluated Shh-induced Hh signaling in Numb CRISPR/Cas9 KO cells. We stimulated cells with Shh and assessed Hh signaling activity by measuring the transcript levels of the Hh target genes, *Gli1* and *Ptch1*. As predicted, Numb loss severely attenuated Hh signaling activity, particularly at the higher doses of Shh. The lower level of Hh signaling was modestly impacted, while the plateau levels of Hh signaling were significantly diminished in Numb KO cells (Fig. 5b, c). To exclude potential off-target effects of CRISPR/Cas9, we infected the Numb KO cells with lentiviruses that express Numb-V5. Restoring Numb expression in Numb KO cells reinstates Shh-induced Hh signaling to levels comparable to those in WT cells (Fig. 5d).

**Figure 5.**
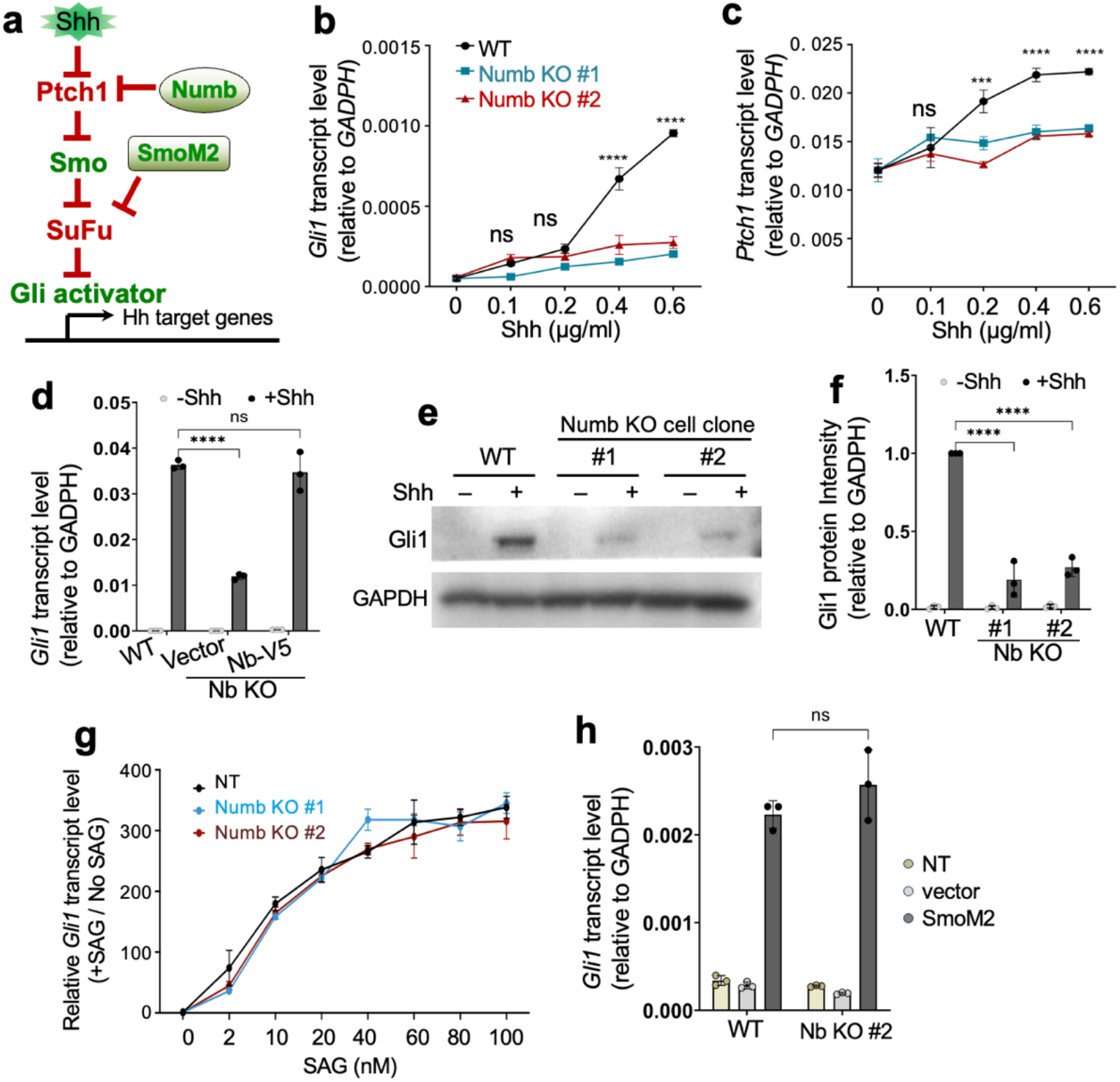
Numb is required for maximal activation of Hh signaling. (**a**) Diagram of the Hh signaling cascade. Numb regulates Hh signaling at the level of Ptch1; SmoM2 actives Hh signaling independent of Shh. **(b-c)** The transcript levels of Hh-target genes, Gli1 and Ptch1, assessed by qPCR. In Numb Ko cells, the transcription of these genes exhibits a significant decrease at high doses of Shh, but remain unaffected at lower doses. (**d**) Hh signaling levels in WT cells and Numb KO cells that express Numb-V5. Numb-V5 reinstates Shh-induced Hh signaling in Numb KO cells. Hh activity was assessed by Gli1 transcript levels. (**e-f**) Western blot analysis and quantification of Gli1 protein in WT and Numb KO cells. Prior to harvesting, cells were stimulated with 1 μg/ml Shh in the low serum culture medium for 24 h. Experiments were repeated four times. (**g**) SAG-induced Hh signaling levels in WT cells and Numb KO cells. Prior to harvesting, cells are treated with the indicated doses of SAG in the low serum culture medium for 24 h. (**h**) WT cells and Numb KO cells were infected with lenticules that express SmoM2. Hh signaling activity was assessed by Gli1 transcript levels. Statistics in (**b**, **c**, **d**, **f**, **g**, **h**): Two-way ANOVA with multiple comparisons (Tukey test). All error bars represent SD. Experiments were repeated four times. ***, p < 0.001; ****, p < 0.0001; ns, not significant.

To confirm the effect of Numb loss on Hh signaling, we silenced Numb expression with an alternative method. We infected cells with lentiviruses that express Numb shRNA and stimulated cells with the highest dose of Shh. Two independent shRNA constructs against Numb attenuated Shh-induced *Gli1* and *Ptch1* transcription to the same extent as Numb CRISPR/Cas9 mediated KO (Supplementary Fig. 9a-c). Further, the attenuated Hh signaling was rescued by the expression of the full length Numb, but not any of the truncated Numb variants that lack either the Ptch1-binding domain (N-terminus and PTB) or the C-terminal domains known to bind to adaptins, components of clathrin-mediated endocytic machinery^42, 43^ (Supplementary Fig. 9d). Thus, both the Ptch1-binding domain and the adaptin-interacting domains are essential for Numb’s positive role in Hh signal transduction.

Numb has been reported to regulate Gli1 protein degradation^44^. We hence tested the impact of Numb loss on the protein levels of Gli1 with Western blot analysis. We found that Gli1 protein levels are significantly reduced in Numb KO cells compared to WT cells (Fig. 5e, f). The reduced Gli1 protein levels are most likely a consequence of attenuated Gli1 transcription.

Given that Numb exerts its regulatory effect at the level of Ptch1, activating the pathway downstream of Ptch1 should turn on Hh signaling in Numb KO cells (Supplementary Fig. 10a-a_3_). To test this, we conducted the following experiments. First, we stimulated cells with SAG, a small chemical agonist of Smo. Across all tested concentrations of SAG, Hh signaling was induced in Numb KO cells to levels comparable to those in WT cells (Fig. 5g). Second, we transfected cells with SmoM2, a constitutively active Smo mutant that triggers Hh signaling independent of Shh^45^. SmoM2 triggered Hh signaling in Numb KO cells to levels comparable to WT cells (Fig. 5h). Third, we knocked down Sufu, a core component of Hh signaling that suppresses the pathway at the level of Gli. Sufu knockdown turned on Hh signaling independently of Shh in Numb KO cells (Supplementary Fig. 10b, c). Finally, we specifically blocked the ciliary activity of PKA, the kinase that exerts its inhibitory roles on Hh signaling in cilia^21, 46, 47^. Expressing a cilium-targeting PKI (a PKA peptide inhibitor tagged to Arl13b-N-RVEP-PR) triggered Hh signaling to a similar magnitude in Numb KO cells and WT cells (Supplementary Fig. 10d). In summary, after Numb depletion, activating Hh signaling downstream of Ptch1 triggers the full activation of Hh signaling, circumventing the inhibitory effects caused by Numb loss.

### Numb loss blocks the activation of Gli transcription factors

We next sought to determine how Numb loss impacts Hh signaling events downstream of Ptch1. We first examined the cilium accumulation of Smo. Interestingly, we found that following Shh stimulation, the intensity of Smo in the cilium of Numb KO cells was comparable to that in WT cells (Supplementary Fig. 11a, b). It is known that Smo may accumulate in the cilium in an inactive conformation^48^. We hence examined the Gli transcription factors downstream of Smo.

At resting state, the transcription of Hh target genes is suppressed by Gli3R, the primary transcriptional repressor. Low levels of Shh stimulation terminate the production of Gli3R, lifting the inhibition on Hh signaling. Conversely, high levels of Shh stimulation activate Gli2, the primary transcriptional activator, resulting in maximal activation of Hh signaling. We first examined the cilium translocation of Gli2, a critical step for Gli2 activation. We found that Shh-induced cilium transport of Gli2 was markedly reduced in Numb KO cells compared with WT cells (Fig. 6a-b). We then analyzed the proteolysis of Gli3, a process responsible for generating Gli3R. The results show that Gli3R production was ceased by Shh stimulation in both WT and Numb KO cells (Fig. 6c-d). These results suggest that without Numb, cells retain the capacity to block the generation of the repressive factor Gli3R, but have deficiencies in activating Gli2.

**Figure 6.**
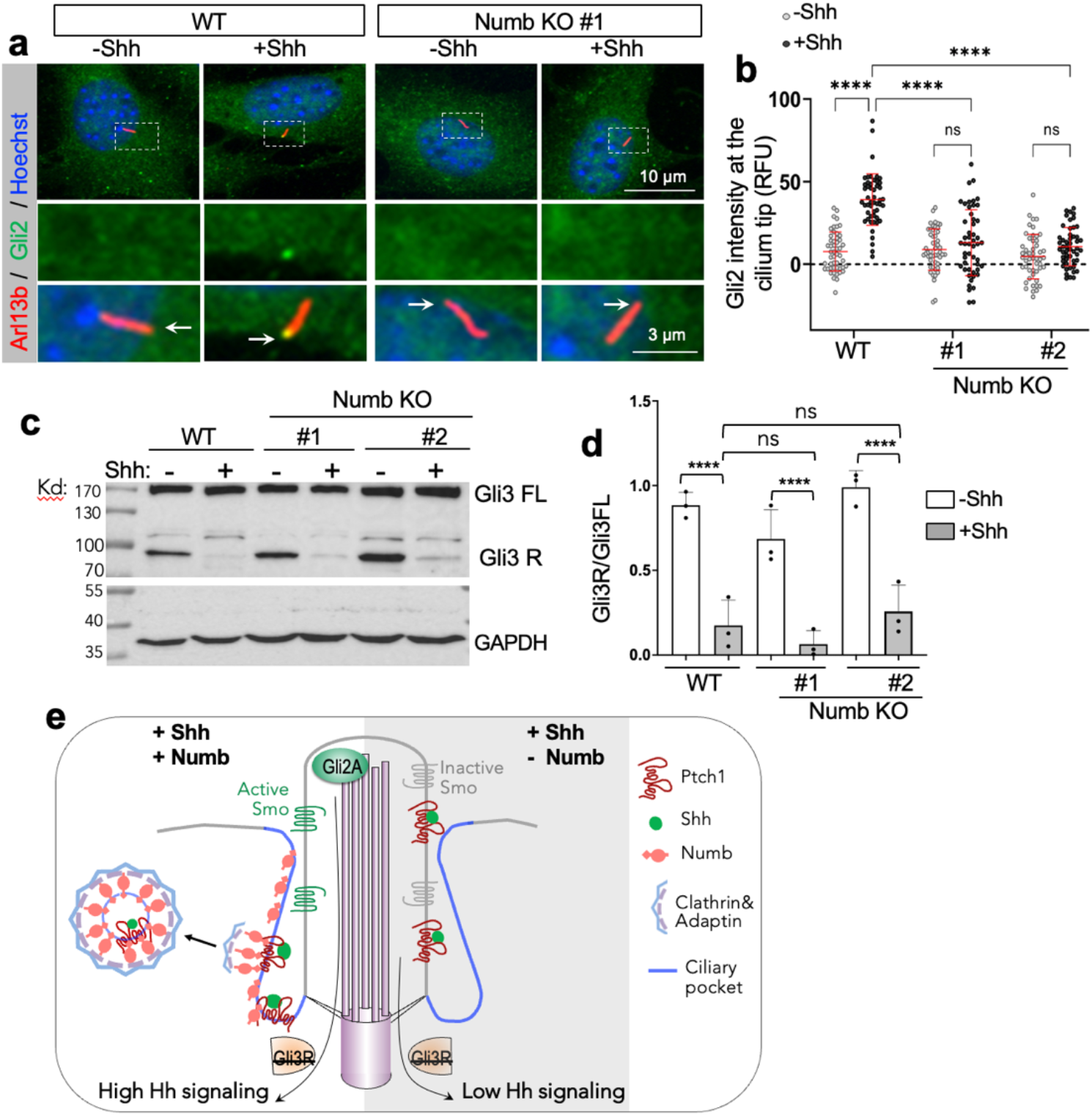
Numb loss blocks the activation of Gli transcription activator. (**a**) Immunofluorescence staining of endogenous Gli2 in the cilia of WT or Numb KO cells. The cells were treated in low serum medium for 24 h, with or without Shh. Enlarged views of highlighted areas are displayed at the bottom. Arrows point to Gli2 fluorescence signal at the cilium tips. (**b**) Quantification of Gli2 fluorescence intensity at the cilium tips. A total of 50 cilia were quantified per condition. RFU, relative fluorescence unit. (**c-d**) Western blot results and quantification of Gli3 processing in WT and Numb KO cells. Cells were stimulated with Shh for 24 h prior to harvest. (**e**) Schematic of Numb’s role during the activation of Hh signaling. Upon Shh binding to Ptch1, Numb in the ciliary pocket (blue lines) recruits Ptch1-Shh complex into clathrin-coated vesicles, thereby facilitating removal of Ptch1 from the cilium. Ptch1’s efficient clearance from the cilium sets the stage for the complete activation of Smo and Gli2 transcription factor. This activation ultimately culminates in the maximal activation of Hh signaling. In the absence of Numb, Ptch1 remains in the cilium even after it binds to Shh. This leads to only partial activation of Smo, which ceases Gli3R production but is insufficient to activate Gli2. As a result, Hh signaling is only moderately activated. Statistics in (**b, d)**: Two-way ANOVA with multiple comparisons (Tukey test). Results from three independent experiments are analyzed. Error bars represent SD. ***, p < 0.001; ****, p < 0.0001; ns, not significant.

The homolog Numblike (Numbl) functions redundantly with Numb under specific circumstances such as neurogenesis in the cerebral cortex^24–26^. We therefore analyzed the localization of Numbl and its role in Hh signaling. We found that Numbl-V5 also localizes to the lower section of the primary cilium. However, unlike Numb, Numbl does not appear as discrete puncta in the cytosol (Supplementary Fig. 12a). This distinction indicates that Numbl may not be involved in endocytosis. We then determined whether Numbl is required for Hh signal transduction. We silenced Numbl with shRNA and found that it does not impact Shh-induced Hh signaling (Supplementary Fig. 12b-c). Further, silencing Numbl in Numb KO cells does not further reduce Shh-induced Hh signaling (Supplementary Fig. 12c). Thus, Numbl is not essential for Hh signaling.

Collectively, our results suggest a molecular model of Numb’s roles at the ciliary pocket in Hh signal transduction (Fig. 6e). As a well-established molecular adaptor for endocytosis, Numb connects the cargo Ptch1 to components of the clathrin-mediated endocytic machinery. Specifically, the N+PTB domain of Numb binds to Ptch1, while its C-terminus interacts with adaptins, thereby incorporating Ptch1 into clathrin-coated vesicles (CCVs). This in turn facilitates Ptch1’s departure from the cilium. While there is a basal level of CCV formation incorporating Ptch1, the binding of Shh to Ptch1 significantly increases the rate of CCV formation, accelerating Ptch1 exit from the cilium. This allows full activation of Smo and Gli2 to induce high levels of Hh signaling. In the absence of Numb, Ptch1 remains in the cilium even after it binds to Shh. The persistent presence of Ptch1 in the cilium impedes the full activation of the signaling steps downstream of Ptch1. Subsequently, cells can cease the generation of Gli3R, but are unable to activate Gli2. As a result, Hh signaling is only moderately activated in absence of Numb.

### Numb is required for high-level Hh responses in spinal cord neural progenitor cells (NPCs)

Hh signaling plays a crucial role in neural tube patterning, where a gradient of Shh ligand secreted by the notochord and floor plate determines distinct neural identities^49, 50^. The identity of each cell fate corresponds to the expression of particular transcription factors within distinct progenitor domains. Along the ventral-dorsal axis, the expression of transcription factors roughly corresponds to the magnitude of Hh signaling. The highest Hh signaling level specifies *FoxA2* expression; high signaling, *Nkx2.2*; medium-to-low signaling, *Olig2* and *Nkx6.1*; and no Hh signaling, *Pax6* (Fig. 7a). This morphogenic activity of Hh ligands can be recapitulated in cultured spinal cord neural progenitor cells (NPCs) derived from embryonic stem (ES) cells^51, 52^. In cultured NPCs, varying levels of Hh signaling elicit the expression of distinct transcription factors, which can be assayed by qPCR or immunostaining.

**Figure 7.**
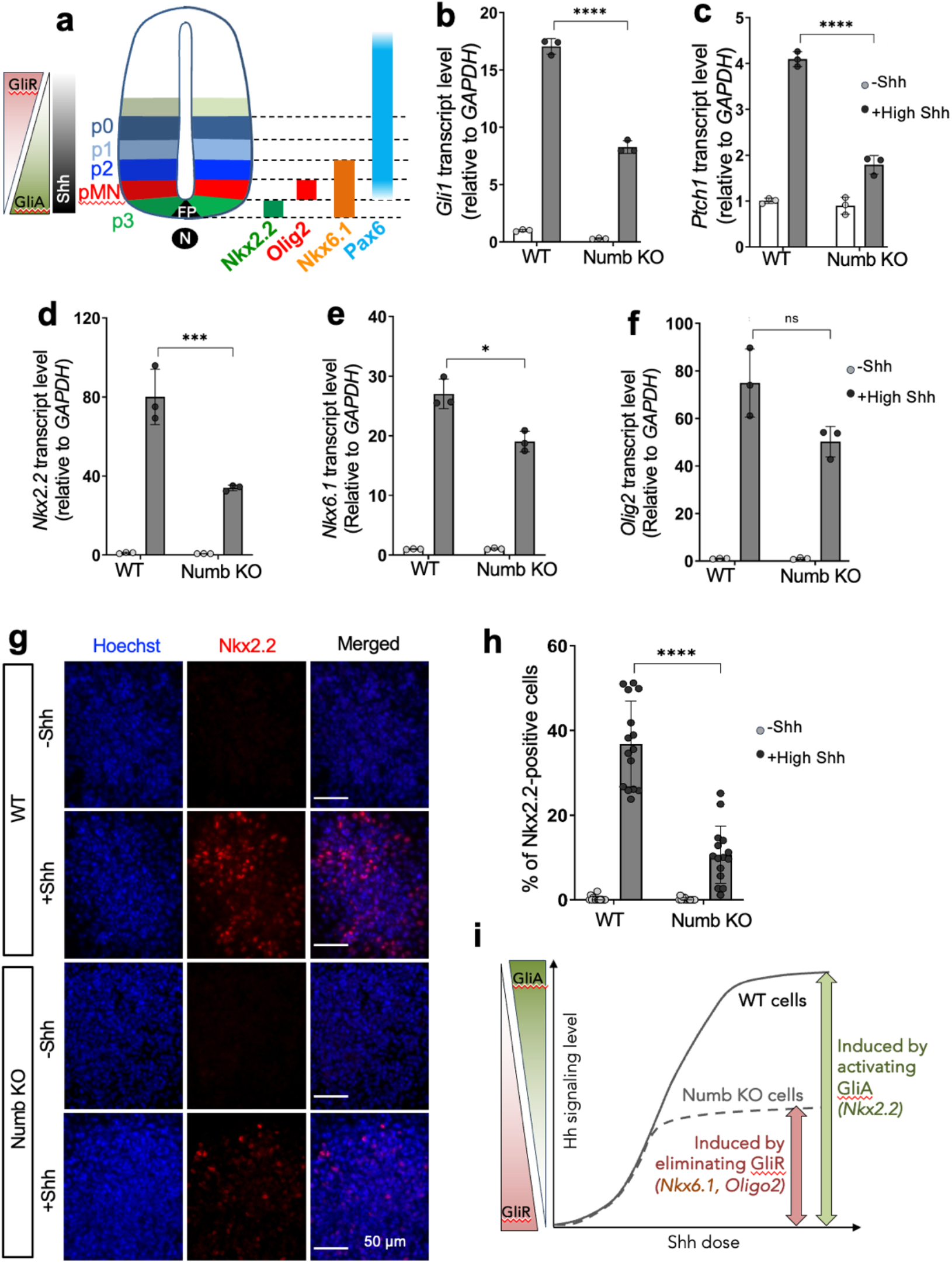
Numb is required for high-level Hh responses in mouse spinal cord NPCs. (**a**) Diagram illustrating embryonic spinal cord progenitors and their associated markers (adapted from Ribes, 2009, *CSH Perspect Biol*). The gradient of Shh along the ventral-to-dorsal axis, ranging from high to low, determines the delineation of progenitor domains. The bars on the right side depict the specific transcription factors in respective domains. N, notochord; FP, floor plate; pMN, motor neuron progenitors; p0, p1, p2, and p3, ventral interneuron progenitors. (**b-c**) The transcript levels of Hh-target genes, *Gli1* and *Ptch1*, in WT and Numb KO cells of mouse spinal cord neural progenitors (NPCs). NPCs underwent differentiation in two conditions: either retinoic acid (RA) (-Shh) or RA with 100nM Shh (+High Shh) for 3 days prior to harvesting. (**d-f**) Transcription levels of specific transcription factors induced by high Shh in WT and Numb KO NPCs. Cells were differentiated in RA-containing medium without or with 100nM Shh for 3 days. Three independent differentiation experiments were performed with similar results. Each condition has three biological replicates. Each dot represents the average of three technical repeats. (**h**) Representative images of NPC differentiation into NKX2.2-positive neural progenitors after 3 days induction. (**i**) The percentage of cells positive for NKX2.2 in WT or Numb KO NPCs. The total of 15 images from different colonies were quantified. Each image represents one NPC colony consisting of approximately 200–300 cells. Each dot represents the data from one image of an NPC colony. Statistics in (**b**-**g, i**): Two-way ANOVA with multiple comparisons (Tukey test). All error bars represent SD. *, p < 0.05; ***, p < 0.001; ****, p < 0.0001; ns, not significant. (**j**) Schematics of Hh signaling and NPC differentiation. Cell fate determination of NPCs in the spinal cord is ultimately determined by the opposing gradient of Gli activator (GliA) and Gli repressor (GliR). High Hh signaling, driven by GliA, induces the differentiation of ventral cell type, such as Nkx2.2-positive cells; whereas medium to low Hh signaling, resulting from the elimination of Gli3R, facilitates the differentiation of dorsal cell types like Olig2- and Nkx6.1-positive cells. In Numb-null ES cells, even with high Hh stimulation, GliA activation is impeded, but cells are still able to curb Gli3R production. Consequently, Numb-null ES cells fail to differentiate into Nkx2.2-positive cells but retain the capacity to differentiate into Olig2- and Nkx6.1-positive NPCs.

To evaluate the role of Numb in NPC differentiation, we employed the same gRNA for NIH3T3 cells and knocked out Numb via CRISPR/Cas9 in mouse ES cells (Supplement Fig. 13a). Since Numb primarily influences the plateau levels of Hh signaling and has minimal effect on the low levels, we exposed cells to high Shh concentrations during their differentiation. As in NIH3T3 cells, Numb loss in NPCs markedly reduced Shh-induced Hh signaling (Fig. 7b, c). Consequently, we observed a substantial reduction in the expression of *Nkx2.2*, in contrast to WT cells. There was moderate impact on the expression of *Nkx6.1*, and no significant effect on *Olig2* (Fig. 7d-f). These results suggest that Numb loss blocks the differentiation of NPCs into Nkx2.2-positive progenitors, the cell identity reliant on high Hh signaling. To further confirm this result, we assayed the differentiation of NPCs into Nkx2.2-positive progenitors via immunostaining with an antibody recognizing Nkx2.2. Following high Shh induction, there is a substantial reduction in the percentage of Numb KO cells positive for Nkx2.2, when compared to WT cells (Fig. 7g-h). In contrast, Numb loss has a moderate or no significant impact on the differentiation of NPCs into Nkx6.1- or Olig2-positive cells (Supplementary Fig. 13 b-d).

Collectively, the results in spinal cord NPCs substantiate our findings in NIH3T3 cells. In the developing spinal cord, along the ventral-dorsal axis, the Shh gradient is converted into opposing gradients of GliA (ventral high, dorsal low) and GliR (dorsal high, ventral low) (Fig. 7a, i). It has been demonstrated in Gli mutant mice that the activation of the ventral marker Nkx2.2 relies on GliA, while the expression of more dorsal markers, Nkx6.1 and Olig2, is curtailed by GliR binding^50, 53^. This is supported by our findings in the cultured NPCs. Our results reveal that in cells depleted of Numb, Gli2 activation is impeded (Fig. 6a), leading to suppression of NPC differentiation into Nkx2.2-positive cells. Meanwhile, Numb-depleted cells manage to eliminate Gli3R, and the remaining moderate levels of Hh signaling is sufficient to induce NPC differentiation into Nkx6.1- and Olig2-positive cells (Fig. 7i).

### Numb is required for Hh signaling and Shh-induced proliferation of GCPs

During cerebellum development, Shh acts as a mitogen to stimulate the proliferation of granule cell precursors (GCPs) in the external granule layer (EGL) from E17.5 to early postnatal development^54–57^. Previous studies have revealed the expression of Numb in the cerebellar GCPs using various techniques, such as RNA-seq^58^, immunostaining^59^, and in situ hybridization^60^ (Supplementary Fig. 15a). We thus determined the subcellular localization of Numb in GCPs. We cultured GCPs obtained from the cerebellum of P7 mouse pups, and infected the GCPs with lentiviruses expressing either Numb-GFP or Numb-mCherry (Fig. 8a). Immunostaining results reveal that both Numb constructs localize to the bottom section of the primary cilium, in addition to its punctate localization in the cytosol. This pattern resembles the localization of Numb observed in NIH3T3 cells (Fig. 3a).

**Figure 8.**
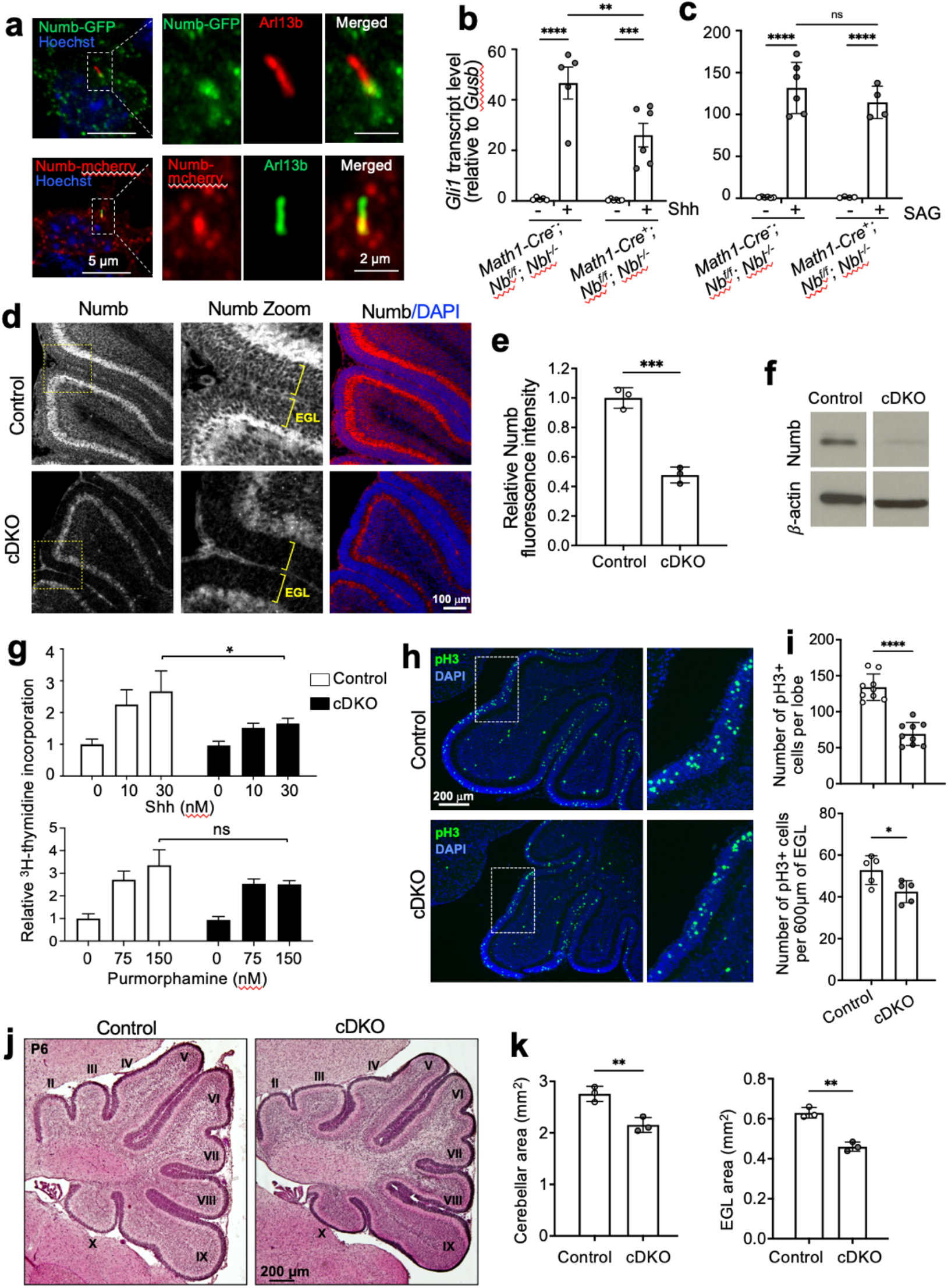
Numb is required for Hh signaling and Shh-induced proliferation in GCPs. (**a**) Numb localizes to the bottom sections of the primary cilium in primary cultured GCPs. GCPs are infected with lentivirus expressing Numb-V5 or Numb-mCherry, and co-stained with ciliary marker Arl13b. **(b-c)** GCPs were isolated from P6 mice and cultured in the absence or presence of (**b**)1 nM Shh or (**c**)150 nM SAG for 20 h. mRNA was isolated from the GCPs and *Gli1* mRNA levels measured by qRT-PCR, relative to *Gusb* mRNA. The deletion of Numb reduces the levels of Gli1 induction by Shh (n ≥ 5 animals per genotype) but not by SAG (n ≥ 4 animals per genotype). *Nb*, *Numb; Nbl, Numblike*. (**d**) Sagittal sections of cerebellum from P6 control and *Numb;Numbl* cDKO mice immunostained for Numb. Numb is most strongly expressed in the EGL and the Purkinje cells. The immunofluorescence intensity of Numb in the EGL of cDKO cerebellum is much lower than that in the control cerebellum. Boxes indicate the zoomed regions. The EGL, external granule layer, is denoted by brackets. (**e**) Quantification of the mean Numb fluorescence intensity in the EGL of control and *Numb;Numbl* cDKO cerebellum. n ≥ 3 cerebella/group. Genotype for control mice: *Math1-Cre^-^;Nb^f/f^;NbL^f/f^*, or *Math1-Cre^-^;Nb^f/f^;NbL^f/-^*, or *Math1-Cre^-^;Nb^f/-^;NbL^f/-^*. Genotype for cDKO mice: *Math1-Cre^+^;Nb^f/f^;Nbl^f/f^*. (**f**) GCPs were isolated from the cerebella of control and Numb;Numbl cDKO mice and the cell lysates analyzed by Western blotting. Numb protein is significantly reduced in GCPs from the cDKO cerebellum. (**g**) GCPs isolated from the cerebellum of P6 control and Numb;Numbl cDKO mice were cultured for 48 h in the presence of Shh or Purmorphamine at the indicated doses. GCP proliferation was measured by the amount of ^3^H-thymidine incorporation. The levels of ^3^H-thymidine incorporation were calculated relative to that in untreated GCPs. n ≥ 3 cerebella per group. Numb loss significantly reduced GCP proliferation in response to the high dose of Shh, but not to Purmorphamine. (**h**) Sagittal sections of cerebellum at the same mediolateral levels from P6 control and *Numb;Numbl* cDKO mice immunostained for the mitotic marker pH3. Boxes indicate the zoomed regions. (**i**) Quantification of the number of pH3-positive cells in the EGL in one lobe of the cerebellum, and pH3-positive cells per 600 μm of the EGL. n ≥ 3 cerebella per group. Deletion of *Numb* and *Numbl* results in fewer proliferating GCPs in the EGL compared to control. (**j**) Sagittal sections of cerebellum at the same mediolateral levels from P6 control and *Numb;Numbl* cDKO mice, stained with hematoxylin and eosin (H & E). Roman numerals indicate the lobules. (**k**) Quantification of the overall cerebellar area and EGL area in P6 mice measured at the same mediolateral level. n = 3 cerebella per group. All Results are shown as mean ± SEM. Statistics in (**e, I, k**): Student’s t-test. Statistics in (**b, c, g**): Two-way ANOVA. **, p < 0.01; ***, p < 0.001, ****; p < 0.0001; ns, not significant.

Given the requirement for Numb in Hh signal transduction in NIH3T3 cells, we hypothesized that Numb may also be required for Hh signaling in GCPs. To test it, we first assessed the effect of Numb loss on Hh signaling in cultured GCPs. We infected primary cultured GCPs with lentiviruses expressing shRNA against Numb. Two independent shRNA effectively silenced Numb expression and significantly diminished Shh-induced Gli1 transcription (Supplementary Fig. 14a-b). The cilium length in Numb knockdown GCPs remains comparable to that in control cells (Supplementary Fig. 14c). We then assessed Shh-induced Ptch1 exit from the cilium. While Shh markedly induced the clearance of Ptch1 from the cilia in control GCPs, this effect was blocked after Numb knockdown (Supplementary Fig. 14d-e). Further, Shh-induced Smo accumulation in the cilium is comparable in GCPs of Numb knockdown and control cells (Supplementary Fig. 14f-g). Hence, Numb positively regulates Hh signaling in GCPs, and exerts similar effect on Ptch1 and Smo as observed in NIH3T3 cells.

To further evaluate Numb’s effect on Hh signaling in GCPs, we generated Numb conditional knockout (*Math1-cre^+^;Numb^f/f^*) in the Numbl-null (*Numbl^-/-^*) background. We used Math1-Cre to mediate depletion of Numb from the GCPs in the developing cerebellum (Fig. 8d-f). In primary cultured GCPs, Shh markedly induced Hh signaling in Numbl-null GCPs; however, removing Numb in Numbl-null background (*Math1-cre^+^;Numb^f/f^;Numbl^-/-^*) significantly reduced Hh signaling (Fig. 8b). Further, we found that SAG-induced Hh signaling levels were comparable in cells with or without Numb (Fig. 8c), corroborating our findings in NIH3T3 cells that Numb exerts its regulatory function at the level of Ptch1.

Next, we determined whether the reduced Hh signaling in GCPs after Numb depletion corresponds to a change in their proliferative response to Shh. We generated mice conditionally lacking Numb and Numbl (Numb;Numbl cDKO) in GCPs using the Math1-Cre driver. Western blot analysis on isolated GCPs showed effective ablation of Numb protein from Numb;Numbl cDKO mice (Fig. 8f). Further, Numb;Numbl cDKO cerebellum exhibits significantly lower Numb immunofluorescence intensity in the EGL (Fig. 8d-e), reflecting effective removal of Numb from GCPs. In primary cultured GCPs, stimulation with Shh for 48 h induced their proliferation in a dose-dependent manner, measured by ^3^H-thymidine incorporation. Numb;Numbl cDKO GCPs had significantly lower proliferation in response to Shh (Fig. 8g). In contrast, cDKO GCPs showed similar proliferation rate to control cells in response to Smo agonist purmorphamine (Fig. 8g). These results align with our findings in NIH3T3 cells that Numb regulates Hh signaling upstream of Smo. We next examined if Numb loss reduced GCP proliferation *in vivo* in the cerebellum. We used the mitotic marker phosphor-histone H3 (pH3) to measure the number of proliferating cells in the EGL. The results show that the EGL of Numb;Numbl cDKO cerebellum contains significantly fewer proliferating cells compared to the EGL of control mice (Fig. 8h-i). In summary, the reduced Hh signaling caused by Numb loss leads to reduced GCP proliferation in the developing cerebellum.

### Numb/Numbl cDKO mice show a decreased GCP population and reduced cerebellar size

The reduced GCP proliferation in EGL may lead to a decrease in GCP population, which usually leads to a reduction in the size of the cerebellum. To test this, we measured the area of the EGL, which is proportional to the size of the GCP population and the overall cerebellar area in the developing cerebellum. We performed the measurement on sagittal sections from P6 cerebellum at the same mediolateral levels. The results show that both the EGL area and the overall cerebellar area are significantly reduced in Numb;Numbl cDKO mice compared to control mice (Fig. 8j-k). Importantly, Numbl single knockout has no significant impact on the overall cerebellar size at P6 (Supplementary Fig. 15b), suggesting that Numb, but not Numbl, is essential for GCP proliferation.

After their proliferative phase, GCPs differentiate into mature granule neurons, exit the EGL and migrate inwards to populate the internal granule layer (IGL). To determine whether the reduced GCP proliferation impacts the overall outcome of granule neurons, we measured the area of the IGL in cerebellar sections at the same mediolateral levels in adult mice. Our results revealed a significant reduction in the IGL area in Numb;Numbl cDKO mice when compared to control mice (Supplementary Fig. 15c-e). Furthermore, the total area of the cerebellum in adult mice was also significantly reduced in Numb;Numbl cDKO mice compared to control mice. Although the IGL area was diminished, the organization of granule neurons did not appear to be affected. Taken together, these results demonstrate that the reduction in GCP proliferation and cerebellar size observed at P6 is not simply a delay in development. Finally, the total body weight of adult Numb;Numbl cDKO mice is comparable to that of control mice (Supplementary Fig. 15f), indicating that the reduced cerebellar size is not due to an overall decrease in the body size of the Numb;Numbl cDKO mice.

In summary, Numb loss diminishes Hh signaling in GCPs, resulting in decreased GCP proliferation and reduced cerebellar size, while the overall organization of the cerebellum remains largely unaffected. Thus, our *in vivo* results in the cerebellum corroborate our findings in NIH3T3 cells and NPCs that Numb is required for the maximal activity of Hh signaling.

## DISCUSSION

### Cilium proteomics with TurboID revealed distinct groups of cilium proteins involved in Shh signaling

In this study, we leveraged TurboID, an engineered biotin ligase, to profile ciliary proteins. We recovered 108 known cilium proteins, including proteins that are well characterized to translocate to the cilium upon Hh activation, such as Smo and Pald1^6, 20^. Our results reveal a distinct group of cilium proteins, and only slightly overlap with results from previous cilium proteomic studies using APEX^19–21^. We captured a large percentage of membrane-associated proteins, whereas previous studies recovered high numbers of ciliary trafficking (IFTs and BBSome) proteins or actin binding proteins. We employed the following strategies to increase the chance of capturing discrete cohorts of protein in the cilium. First, we leveraged TurboID, a proximity labeling enzyme with distinct mechanisms from APEX. TurboID is derived from the biotin ligase BirA^61^, and mediates biotin tagging to primary amines, such as the lysine sidechain of target proteins. In contrast, APEX is a derivative of ascorbate peroxidase that catalyzes the tagging of biotin-phenol to electron-dense residues such as tyrosine^62^. These distinct labeling mechanisms allow them to target different cohorts of proteins. Second, we targeted TurboID to the juxtamembrane domain by fusing it to truncated Arl13b. This variant of Arl13b has a small size (90aa), is associated to the membrane via lipid-link, and Arl13b has known functional interactions with Hh signaling components. These factors maximized our chance of capturing signal transducers that are recruited to the juxtamembrane region during signaling transduction.

### Numb regulates protein levels in the cilia via mediating endocytosis at the ciliary pocket

Previous research has associated the ciliary pocket with endocytosis of ciliary proteins^36, 37, 63^. However, there has been a lack of study elucidating how this is achieved at the molecular level. To date, no specific molecules have been directly demonstrated to undergo endocytosis from the ciliary pocket. In this study, we identified Numb as part of an endocytic machinery at the ciliary pocket that plays an important role in the control of Ptch1 protein levels in the cilium. We discovered Numb from a cilium proteomic study, and confirmed its localization at the ciliary pocket via expansion microscopy and immuno-electron microscopy (EM) (Fig. 3a-c). We found that Numb interacts with Ptch1 via the Numb-PTB domain and N-terminus. The C-terminus of Numb is known to interact with several components of the endocytic and vesicle sorting machinery, such as α-adaptin and EHD family members^42, 43, 64–66^. Through these molecular interactions, Numb incorporates Ptch1 into clathrin-coated vesicles (CCVs), thereby facilitating Ptch1 departure from the cilium (Fig. 3d-i). This Numb-mediated endocytosis of Ptch1 may also occur at the resting stage, but this process is significantly enhanced upon the binding of Shh to Ptch1 (Fig. 3h-I). It is possible that upon binding to Shh, Ptch1 adopts a conformation that promotes its interaction with Numb. This enhanced interaction could facilitate the efficient incorporation of Ptch1 into CCVs, thereby promoting its internalization from the ciliary pocket. The clearance of Ptch1 from the cilium is essential for the maximal activation of the Hh pathway (Fig. 5-6). Broadly, as an endocytic adaptor, Numb has been reported to mediate the internalization of multiple cell surface receptors in non-ciliary contexts^31, 32, 59, 67^. It is intriguing to speculate that the Numb-clathrin machinery could potentially serve as a universal mechanism to modulate the abundance of diverse proteins in the cilium.

Notably, immunostaining with anti-Numb antibodies did not yield detectable signals for endogenous Numb in the ciliary pocket. This could potentially be attributed to the low levels of Numb in the ciliary pocket. Given its role as an endocytic adaptor protein, it is conceivable that Numb’s presence in the ciliary pocket is transient, resulting in low levels that fall below the detection threshold of the Numb antibody. A similar phenomenon has been noted with other ciliary proteins, such as PKA. Multiple PKA subunits were detected in the cilium in mass spectrometry studies^21^. Further, when PKA subunits were fused to EGFP, their presence was observed in the cilium^21, 68^. However, efforts to detect endogenous PKA in the cilium in WT cells using immunostaining methods have not yielded successful results.

### Numb positively regulates Hh signaling

We found that Numb positively regulates Hh signaling by facilitating Ptch1 exiting from the cilium upon Shh stimulation (Fig. 5). Our findings in NIH3T3 cells are corroborated by results in NPCs and GCPs (Fig. 7-8). Our results differ from a previous report showing that Numb negatively regulates Hh signaling by facilitating the degradation of Gli1^44^. Multiple lines of evidence support the idea that our results reveal the physiological role of Numb in Hh signaling.

First, most experiments in the previous study^44^ were conducted in non-ciliated cell lines, and hence may not be relevant to the physiological process of Hh signal transduction in the vertebrate system. Second, we found that diminished Hh signaling is rescued in Numb KO cells by: 1) restoring Numb expression, 2) stimulating cells with a Smo agonist; 3) expressing a constitutively active Smo mutant, 4) suppressing ciliary PKA with PKI, and 5) knockdown of Sufu. These results show that Numb regulates Hh signaling upstream of Smo, consistent with our identified roles of Numb in Ptch1 internalization from the ciliary pocket. Third, in the developing cerebellum where Hh signaling plays a mitogenic role, genetic depletion of Numb leads to reduced GCP proliferation. This result provides strong evidence that Numb acts as a positive regulator in Hh signaling under physiological conditions. Finally, we found that ablation of Numb markedly reduced Gli1 protein levels, most likely as the consequence of the reduced Gli1 transcription. Gli1, as a target gene of the Hh pathway and an amplifier of Hh signaling, is induced exclusively after Hh signaling is activated. Gli1 protein levels in resting stage remain undetectable. Hence, the scenario described in the previous study may potentially be relevant to special circumstances, such as certain cancer cells where Hh signaling is triggered by mutations downstream of Ptch1. However, it may not be applicable to the physiological process of Hh activation.

Our discovery in NIH3T3 cells is corroborated by our analysis in the cerebellum of Numb;Numbl cDKO mice. We found that GCP proliferation, a Hh-dependent process, is markedly reduced in Numb;Numbl cDKO mice, leading to a reduction in EGL area (IGL in adult cerebellum) and in the overall size of the cerebellum (Fig. 8, Supplementary Fig. 15). In the developing cerebellum, Numb regulates granule cell maturation and Purkinje cell maturation^69^ and is required for BDNF-mediated migration of GCPs after they exit the cell cycle^59^. Consistent with a role for Numb in granule cell maturation and proliferation, conditional removal of Numb with En2-Cre, a Cre line that ablates Numb from the entire hindbrain at early developmental stages, results in a smaller cerebellum^69^. Our findings in NIH3T3 cells also align with the results in induced differentiation of spinal cord NPCs. In Numb-null NIH3T3 cells, Shh stimulation inhibits the generation of Gli3R but does not activate Gli2. Consequently, Numb loss attenuates the plateau level of Hh signaling and has a minimal impact on low levels of Hh signaling. Consistent with this result, the absence of Numb hinders the differentiation of NPCs into cell fate dependent on high Hh signaling (Nkx2.2-positive), while showing moderate to no impact on cell fates reliant on low Hh signaling (Nkx6.1- or Olig2-positive). Notably, our results in NPCs align with the reported phenotype in the spinal cord of Numb cKO;Numblike KO mice, where Olig2-positive progenitor cells appear to be correctly specified^25^.

### Numb is required for maximal activation of Hh signaling

Our results show that in cells lacking Numb, Ptch1 remains in the cilia even after binding to Shh. This permits a moderate level of Hh signaling activation but blocks the attainment of maximal Hh signaling levels. These results are in line with several previous studies, all highlighting that when Ptch1 clearance from the cilium is hindered by various factors including Ptch1 mutations, the maximal activation of Hh signaling is compromised^16, 70, 71^. The straightforward explanation for these observations is that the Ptch1-free ciliary environment is needed for the maximal activation of Hh signaling. Ptch1 is known to suppress Hh signaling by preventing the access of Smo to free cholesterols in the cilium, a step required for Smo activation. The binding of Shh to Ptch1 blocks Ptch1 activity as a cholesterol transporter^8–12^; however, when Ptch1-Shh complex persists in the cilium, the ciliary environment is not conducive to optimal activation of Smo. Therefore, in Numb-loss cells, even when Smo is accumulated in the cilium, Smo is not fully activated. The ciliary Smo may adopt an inactive conformation similar to that induced by cyclopamine^48^; alternatively, among all ciliary Smo proteins, only a subpopulation is activated. Under either condition, the downstream signaling is activated sufficiently to cease Gli3R production but not robustly enough to enable Gli2 activation (Fig. 6). This observation is further substantiated by our fundings in NPCs, where Numb loss allows cell differentiation into Olig2- and Nkx6.1-positive progenitors that rely on medium-to-low Hh activity, but blocks differentiation into cell fates that require high Hh signaling, such as Nkx2.2-positive progenitors.

In a previous study by Yue et al.^16^, Ptch1 levels at the cell surface were regulated by Smurf-mediated ubiquitination and the ensuing degradation in the endosome. Although an increase in Ptch1 levels in the cilium was noted after Smurf knockdown, this may not be the direct consequence of Smurf activity in the cilium, as there is no established evidence of Smurf localization to the cilium. The observed increase in ciliary Ptch1 levels in Smurf-deficient cells could potentially result from the overall elevated Ptch1 levels in the cytoplasmic membrane. Nevertheless, consistent with our observations in Numb-null cells, Ptch1 persists in the cilium even after its binding to Shh in Smurf-deficient cells, leading to attenuated Hh signaling. The Smurf-mediated Ptch1 degradation may operate in conjunction with our current model of Ptch1 exit from the cilium: Numb facilitates the endocytosis of Ptch1 at the ciliary pocket by incorporating Ptch1 into CCVs, and the fate of endocytosed Ptch1 is determined by its ubiquitination status. Collectively, the findings from both Yue et al. and our study underscore the critical significance of clearing Ptch1 from the cilia for the maximal activation of Hh signaling. Our discovery of Numb as a positive regulator of Hh signaling expands our understanding of Hh signaling, and reveals an endocytosis machinery at the ciliary pocket that could play crucial roles in the regulation of ciliary protein levels.

---

**Supplementary Figure 1.**
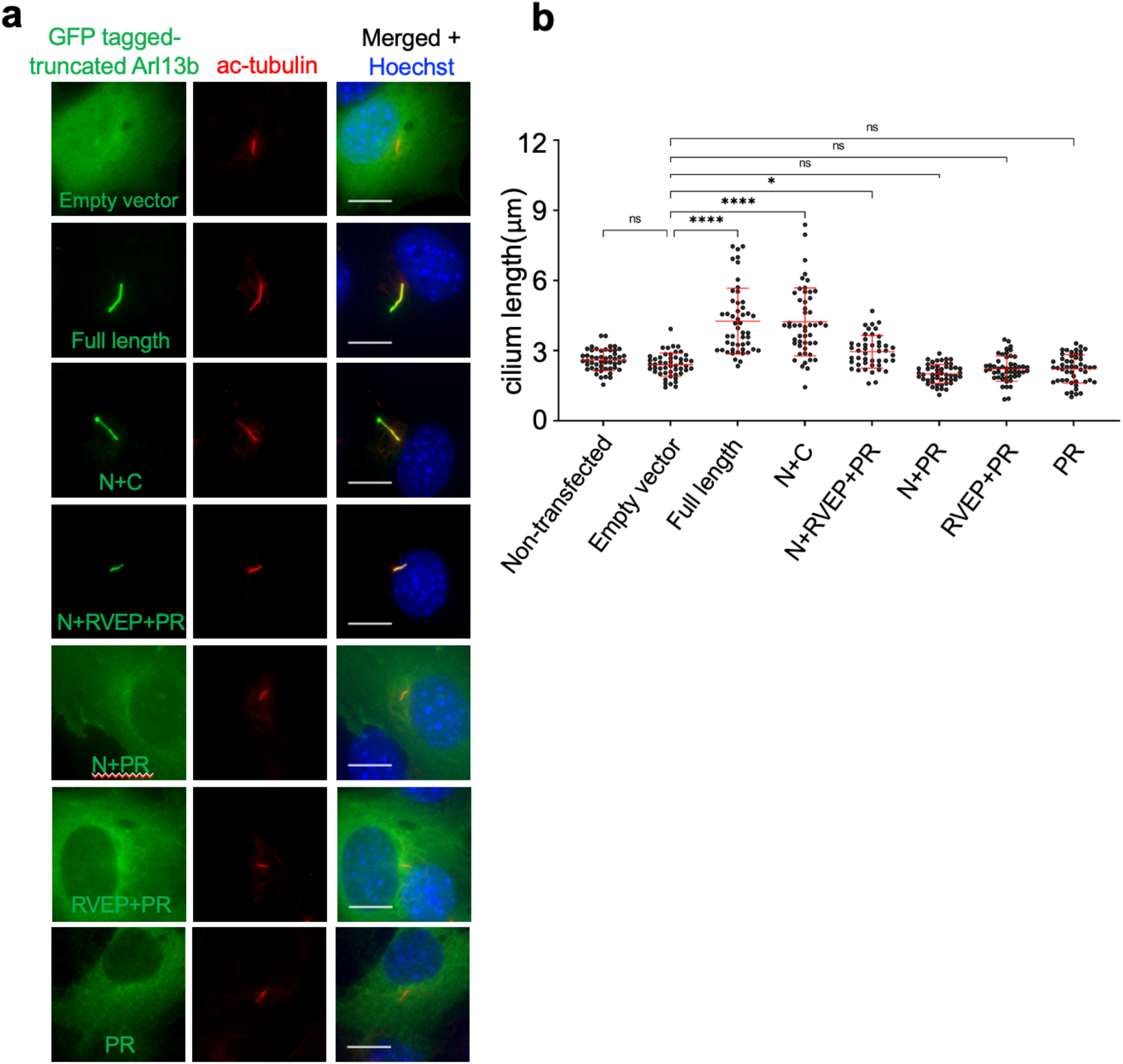
Truncated Arl13b localizes to the cilium with minimal impact on cilium length in transiently transfected cells. (**a**) GFP-tagged truncated Arl13b constructs were transiently expressed in NIH3T3 cells. Cells were immunostained with an antibody against GFP and the cilium marker acetylated-tubulin. Scale bar, 10 µm. (**b**) Quantification of the cilium length in cells transiently transfected with truncated Arl13b constructs. Note that in stable cell lines in Fig S2A where the expression levels are much lower, N+RVEP+PR has no significant impact on cilium length. Data are shown as mean ± SD. A total of 60 cilia were quantified per condition. One-way ANOVA with multiple comparisons (Tukey test). ****, p < 0.0001; *, p < 0.05; ns, not significant. Scale bars, 10 µm in all panels.

**Supplementary Figure 2.**
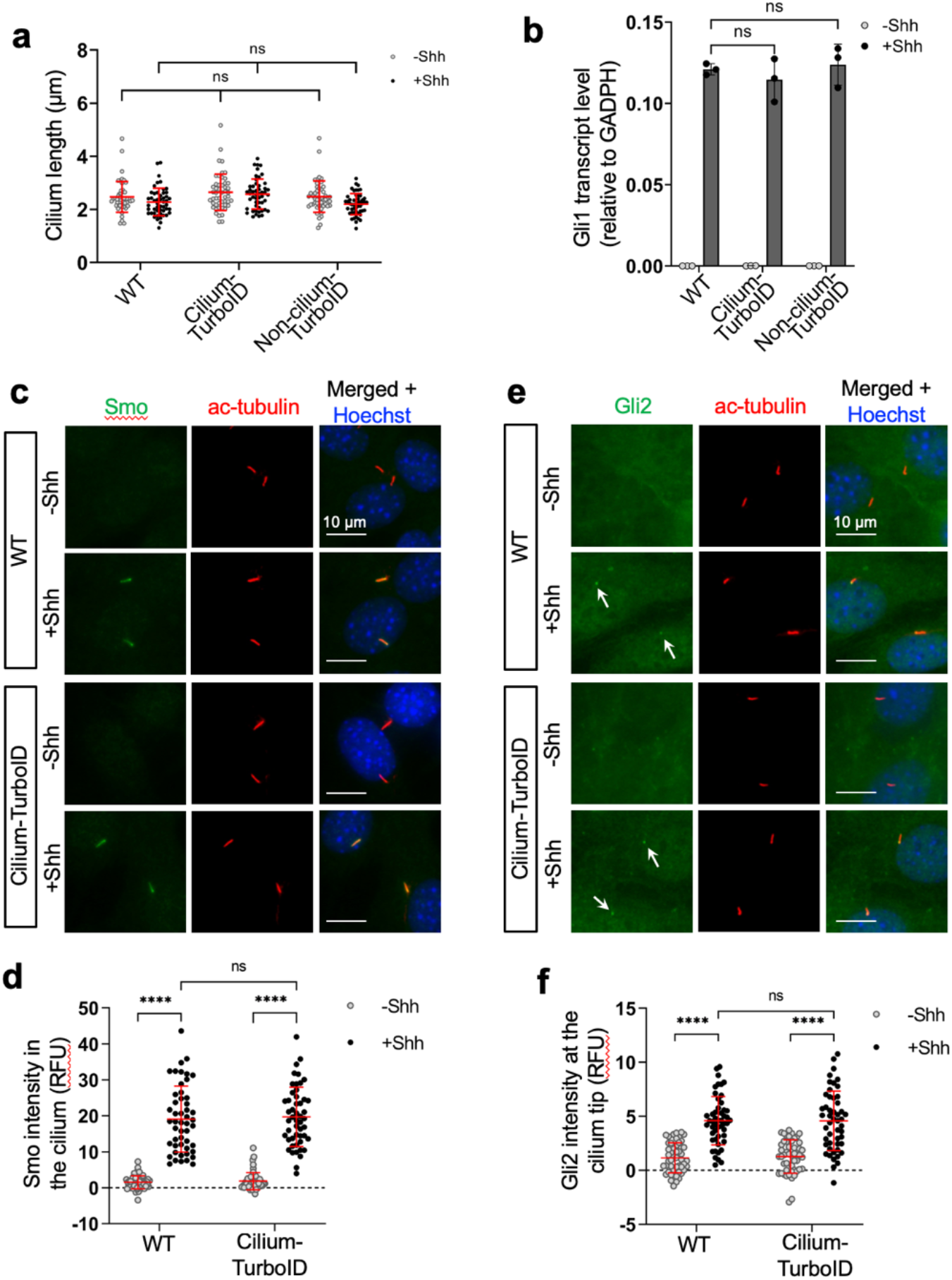
Cilium-TurboID in stable cell lines does not impact cilium morphology and Hh signal transduction. (**a**) Quantification of the cilium length in WT (parental NIH3T3 cells), Cilium-TurboID and Non-cilium-TurboID stable cell lines. A total of 50 cilia were quantified per condition. (**b**) Cells were stimulated with Shh for 24h, and Hh signaling activity was evaluated by qPCR measuring Gli1 transcript levels. Data are shown as relative transcript levels normalized to GAPDH (n = 4). (**c**, **e**) Immunofluorescence of Smo and Gli2 in WT and Cilium-TurboID stable cell lines. Cells were serum-starved with or without Shh in the culture medium for 24 h. Arrows in (**d**) point to Gli2 fluorescence signal at cilia. (**d**, **f**) Quantification of Smo and Gli2 fluorescence intensity in the cilium. Data are shown as mean ± SD. A total of 50 cilia were quantified per condition. RFU, relative fluorescence unit. Statistics in (**a**, **b**, **d** and **f**): Two-way ANOVA with multiple comparisons (Tukey test), ****, p < 0.0001. All error bars represent SD; ns, not significant. Scale bars, 10 µm in all panels.

**Supplementary Figure 3.**
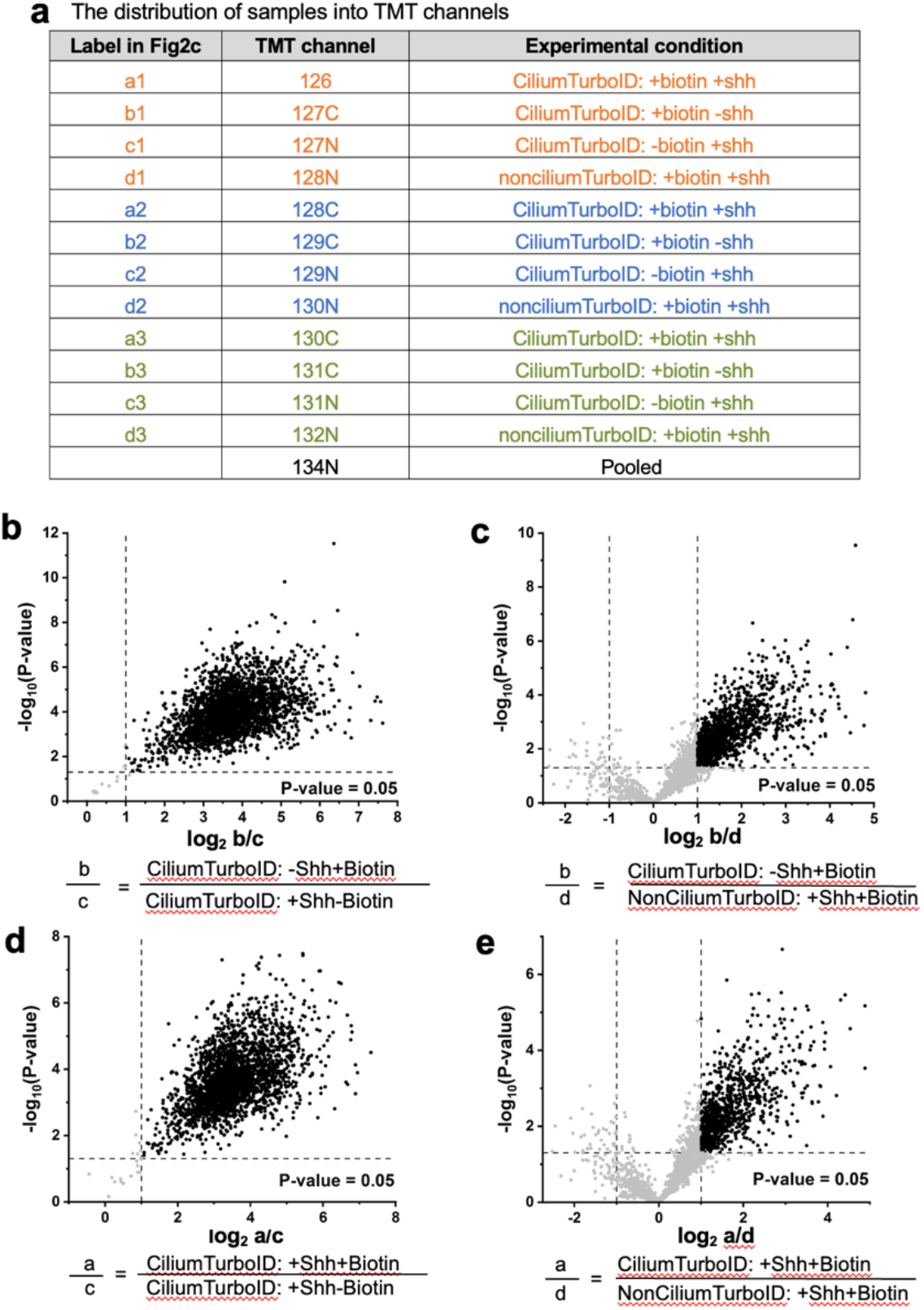
Analysis of mass spectrometry results. (**a**) The distribution of samples in the channels of TMTpro 16plex label reagent set. (**b-e**) Analysis was performed on 2,880 proteins that have more than 7 unique peptides. Volcano plots were generated by plotting the log2 TMT ratio of the indicated conditions against the negative log10 p-value of Student’s t-test of the corresponding conditions. The P values were adjusted for multiple testing via the Benjamini-Hochberg procedure. In each plot, dashed lines indicate the cutoffs of p < 0.05 and TMT ratio > 2 or -2. Proteins that meet the cutoffs are represented by black dots. The proteins that meet the cutoffs were further analyzed as follows. (1) The intersection of proteins in (**b**) (2643 candidates) and in (**c**) (1225 candidates) are defined as ciliary proteins without Shh stimulation; 788 proteins are identified in this category (Table S1). (2) The intersection of proteins in (**d**) (2636 candidates) and in (**e**) (890 candidates) are defined as ciliary proteins with Shh stimulation; 574 proteins are identified in this category (Table S1). (3) The union of candidates identified in step (1) and (2) are defined as ciliary candidates. 800 proteins are identified in this category (Table S2). These 800 proteins are plotted in Fig. 2c and Fig. 2e.

**Supplementary Figure 4.**
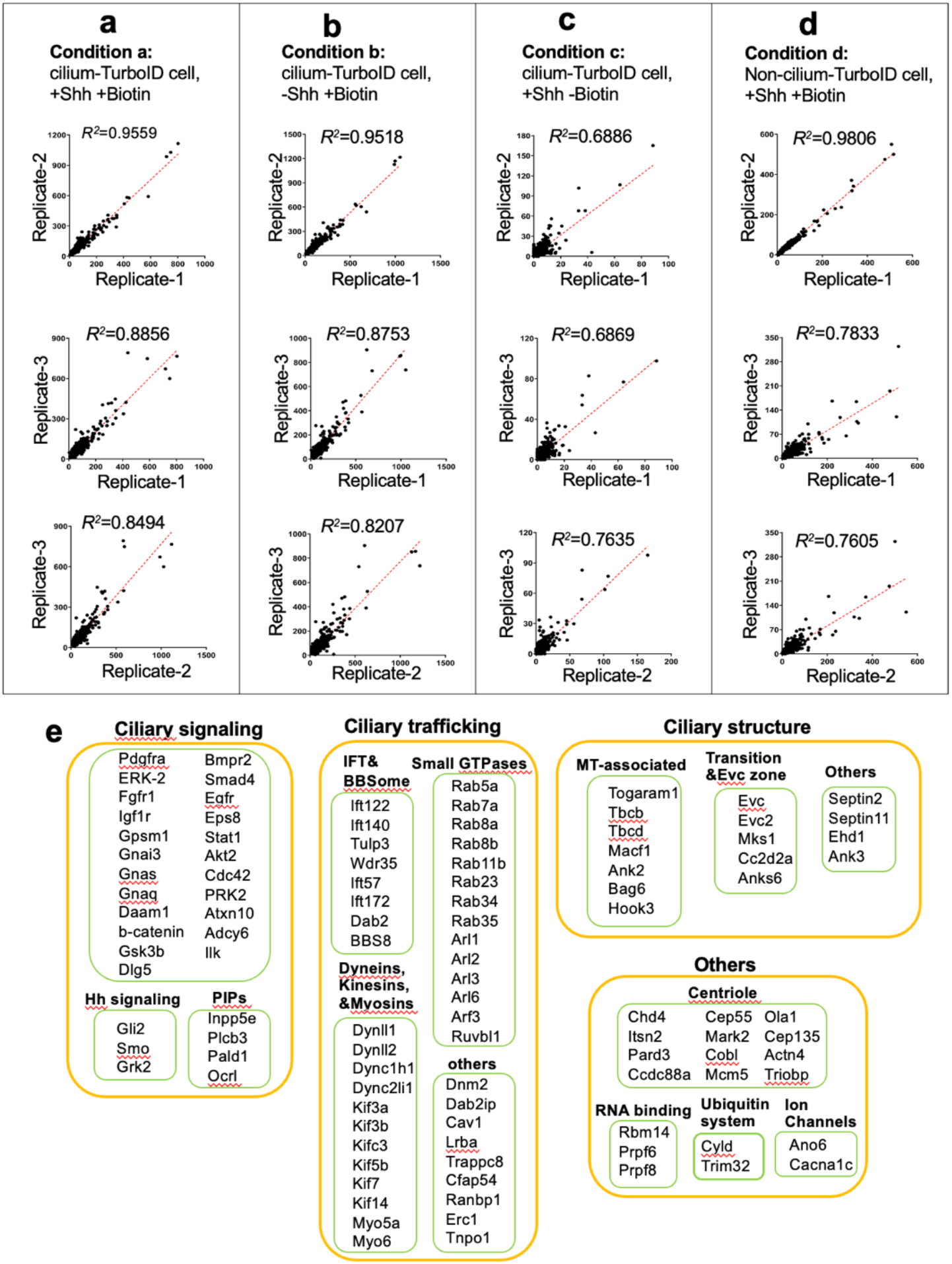
Correlation of biological replicates in mass spectrometry. (**a**) Correlation of biological replicates for condition a: cilium-TurboID cells labeled with biotin and stimulated with Shh. The averaged R-squared is 0.8969. (**b**) Correlation of biological replicates for condition b: cilium-TurboID cells labeled with biotin without Shh. The averaged R-squared is 0.8826. (**c**) Correlation of biological replicates for condition c: cilium-TurboID cells stimulated with Shh Without biotin labeling. The averaged R-squared is 0.7130. (**d**) Correlation of biological replicates for condition d: non-cilium-TurboID cells labeled with biotin and stimulated with Shh. The averaged R-squared is 0.8415. (**e**) The previously reported ciliary candidates recovered in this study are listed and grouped into functional categories.

**Supplementary Figure 5.**
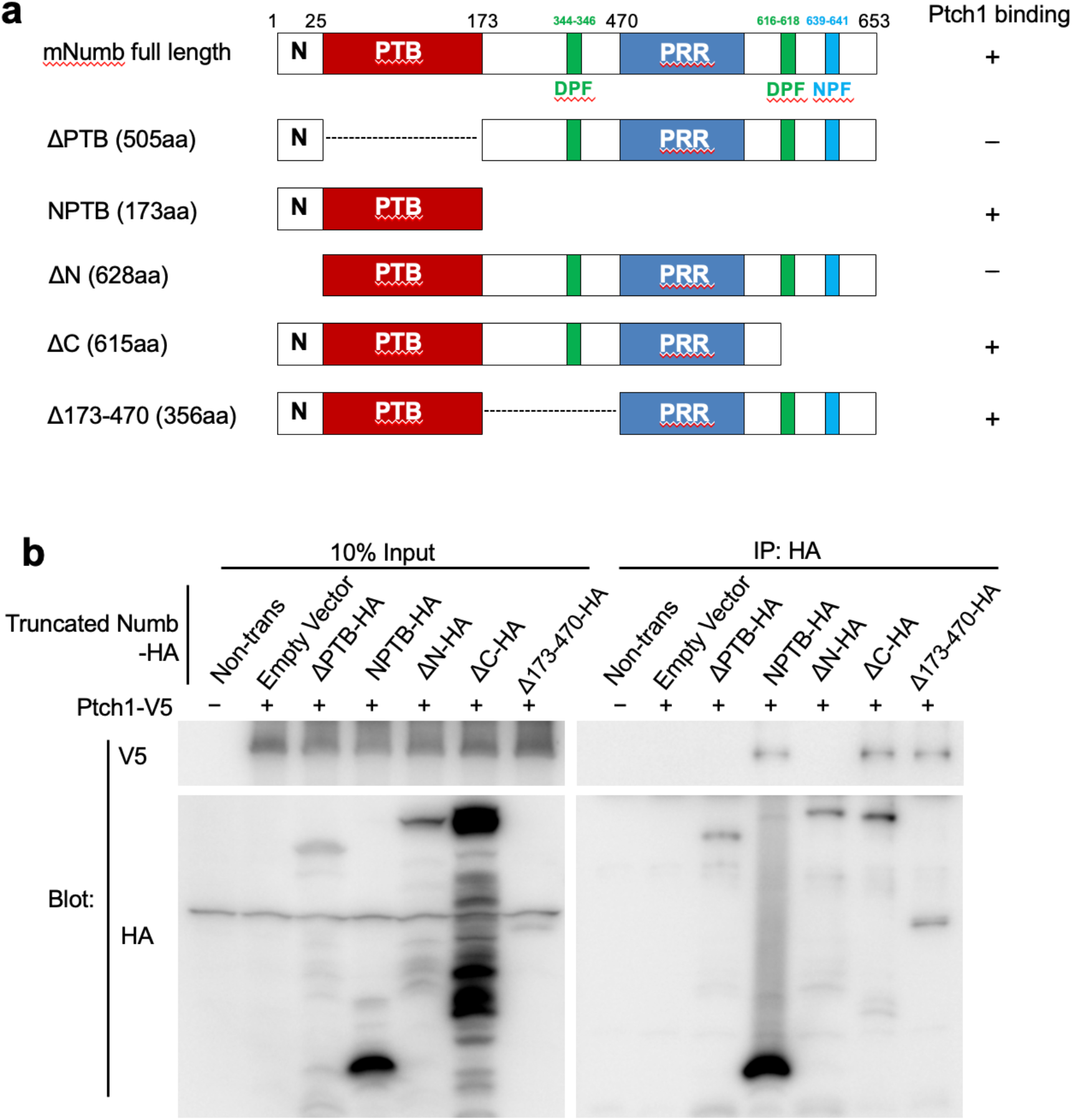
Numb binds to Ptch1 via Numb-N and PTB domain. (**a**) Schematic representation of Numb protein structure and truncated Numb variants used to map the interaction domain with Ptch1. (**b**) Ptch1-V5 and HA tagged-Numb-truncated proteins were expressed in 293T cells. Co-immunoprecipitated was done with an anti-HA antibody. Deletion of either the N or PTB domain disrupts the interaction between Numb and Ptch1, while the N+PTB domain itself is sufficient to mediate the binding with Ptch1-V5.

**Supplementary Figure 6.**
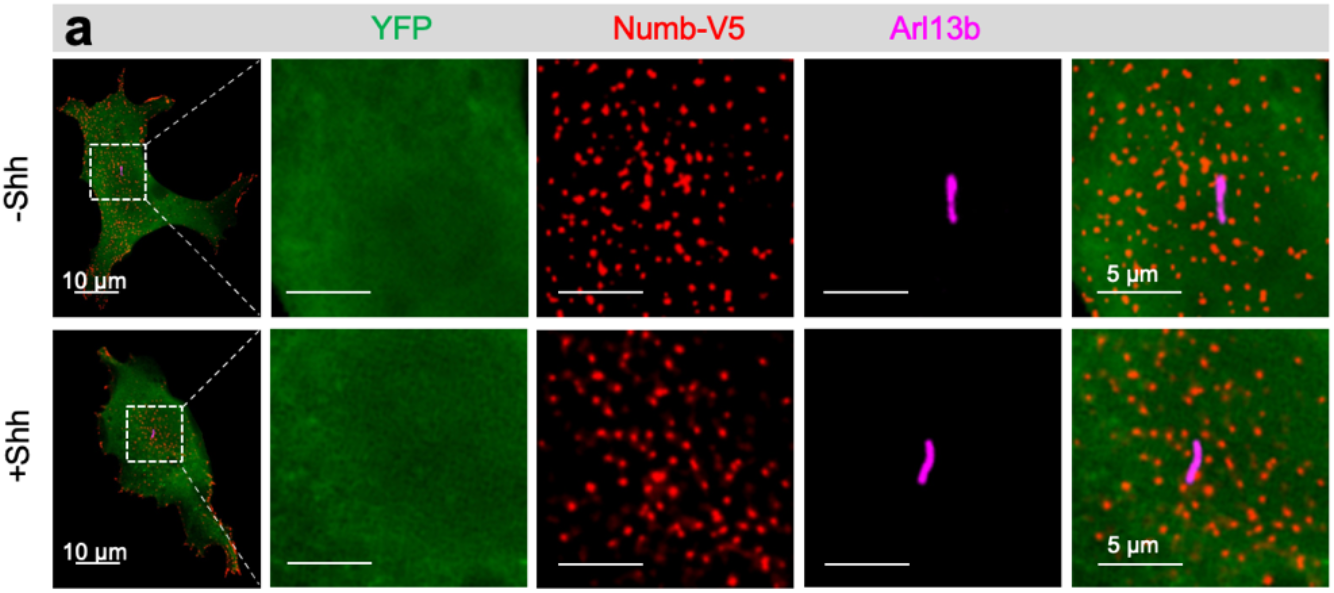
YFP control does not show any co-localization with Numb. (**a**) As a control for Figure 3h, YFP protein was expressed in NIH3T3 cells, together with Numb-V5. Cells were treated with Shh or vehicle for 30 min before fixation and staining. YFP control does not show any co-localization with Numb.

**Supplementary Figure 7.**
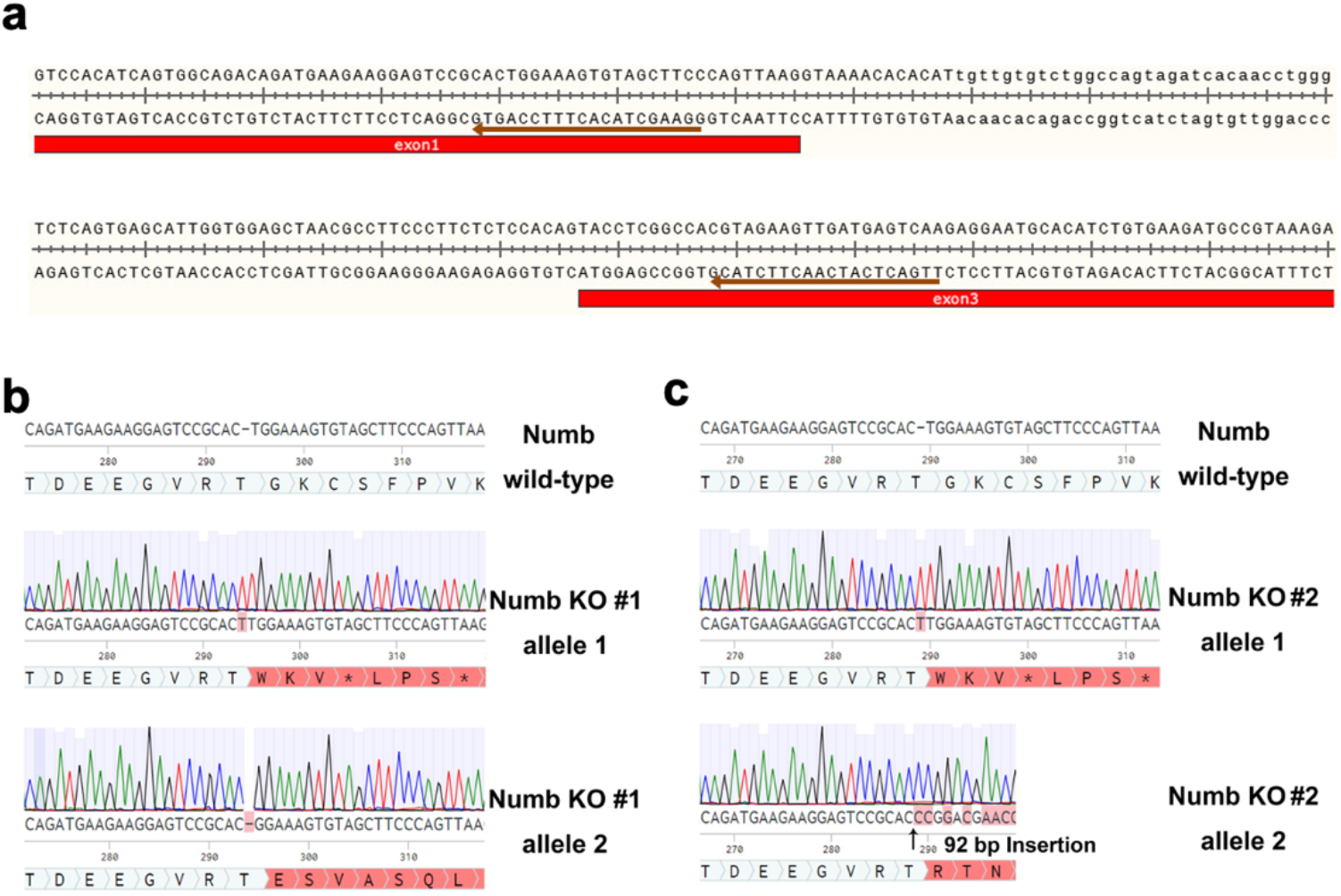
Design of Numb CRISPR/Cas9 and identification of genomic mutations in individual cell clones. (**a**) Two guide RNAs were designed to target exon 1 and 3 of mouse Numb. Brown arrows indicate the sequence of gRNAs. (**b-c**) The gRNA targeting region in the mouse genome was amplified by genomic PCR, ligated into TOPO vector, and transfected into chemically competent cells. 20 bacterial colonies of each cell clones were randomly picked and sequenced. The sequences are aligned with the Numb WT gene sequence from M. musculus. In Numb KO cell clone #1, a single-base-pair insertion and single-base-pair deletion were present. In Numb KO cell clone #2, a single-base-pair insertion and 92bp insertion were present. All mutations cause biallelic frameshift mutations that eventually lead to nonsense-mediated decay (NMD) of the mutant mRNA.

**Supplementary Figure 8.**
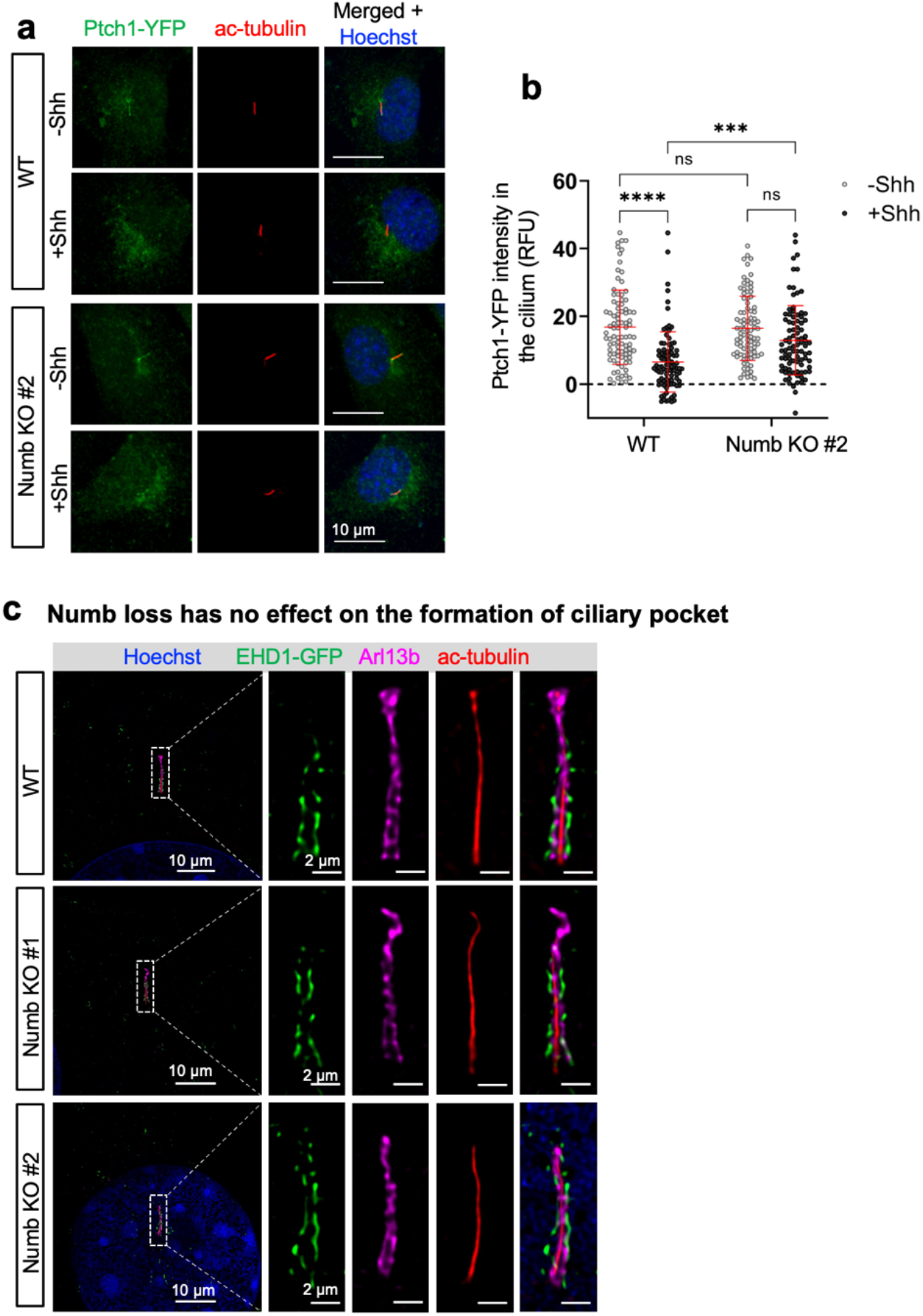
Numb is required for Shh-induced Ptch1-YFP exit from the cilium. (**a**) Representative images of Ptch1-YFP in the cilium before and after Shh treatment. NIH3T3 cells were infected with Ptch-YFP lentivirus and treated with Shh or vehicle for 30 min before fixation and staining. Cilia are labeled with acetylated-tubulin (red). (**b**) Quantification of Ptch1-YFP intensity in the cilia. n = 60-70 cells per condition. Two-way ANOVA with multiple comparisons (Tukey test). All error bars represent SD. ***, p < 0.001; ****, p < 0.0001. ns, not significant. (**c**) Numb loss has no effect on the formation of ciliary pocket. Representative images of the morphology of ciliary pocket in WT and Numb KO cells. Cells were transfected with EHD1-GFP, underwent expansion microscopy, and imaged with Airyscan microscopy. The image shown is from a single focal plane highlighting the ciliary pocket. The cilium axoneme is labeled with acetylated (ac)-tubulin (red); cilium membrane is labeled with Arl13b (Magenta); ciliary pocket is highlighted by EHD1 (green).

**Supplementary Figure 9.**
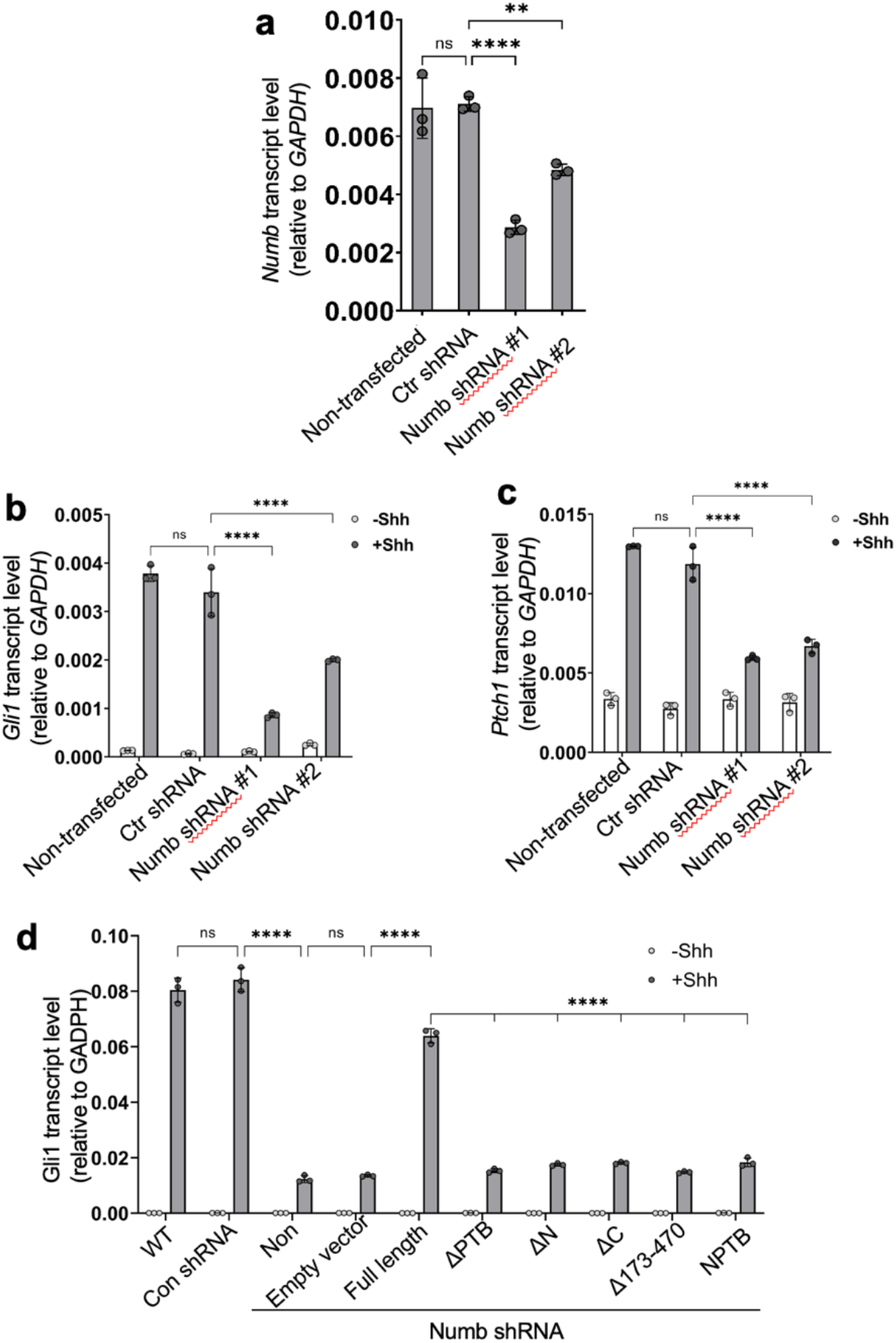
Numb shRNA impairs activation of Hh signaling. (**a**) *Numb* shRNA significantly reduced *Numb* gene expression. *Numb* mRNA levels were measured by qPCR. Control (Ctr) shRNA is a non-mammalian targeting shRNA. (**b-c**) 3 days after lentivirus-mediated transfection of *Numb* shRNA, cells were serum-starved for 24 h with or without Shh. Hh signaling activity was evaluated via qPCR measuring transcript levels of *Gli1* and *Ptch1*. *Numb* knockdown reduced Shh-induced Hh target gene transcription. (**d**) The ShRNA-resistant Numb and Numb truncated variants were expressed in cells of Numb knockdown. The full-length Numb protein, but not any of the truncated variants rescued Hh signal transduction. Hh signaling was assessed by the levels of Gli1 transcription. Statistics in (**a**): One-way ANOVA with multiple comparisons (Tukey test). (**b**-**d**): Two-way ANOVA with multiple comparisons (Tukey test). All error bars represent SD, n=3. ****, p < 0.0001; ***, p < 0.001; **, p < 0.01; *, p < 0.05. ns, not significant.

**Supplementary Figure 10.**
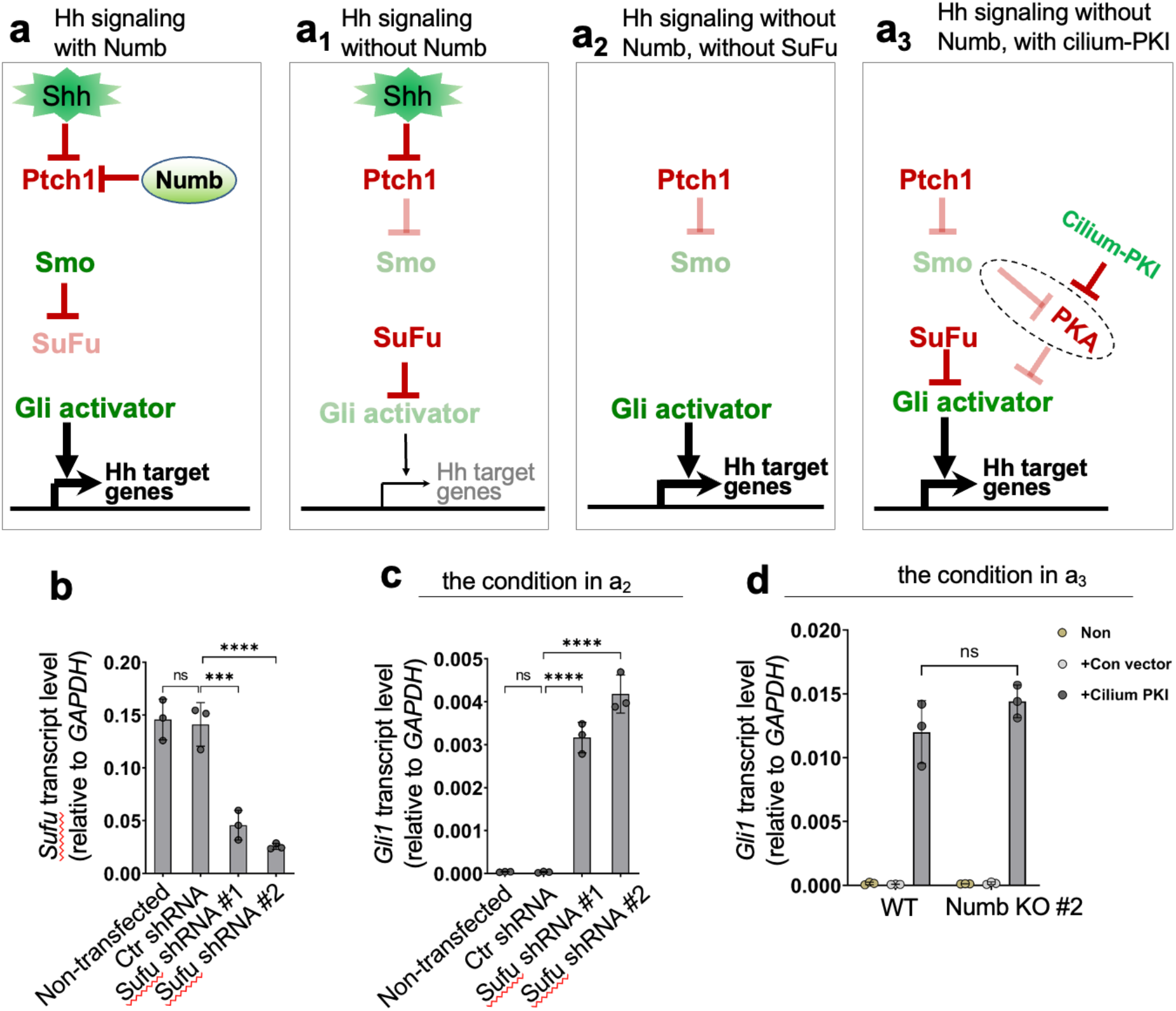
Activating signaling components downstream of Ptch1 turns on Hh signaling in Numb KO cells. (**a-a_3_**) The schematic of Hh signaling in various scenarios. (**a**) Numb enhances Hh signaling at the level of Ptch1; (**a_1_**) Numb loss attenuates the Hh signaling activity; (**a_2_**) SuFu knockdown turns on Hh signaling independent of Shh in Numb knockout (KO) cells; (**a_3_**) Inhibiting ciliary PKA activity turns on Hh signaling independent of Shh in Numb KO cells. (**b**) Sufu shRNAs significantly reduced Sufu gene expression in Numb KO cells. Sufu mRNA levels were measured by qPCR. (**c**) In Numb KO cells, Hh signaling is turned on by Sufu knockdown. Hh signaling activity is assessed by Gli1 transcript levels. This condition corresponds to the scenario **a_2_**. (**d**) PKI fused with Arl13b-N-RVEP-PR (Cilium-PKI) was expressed in cells via lentiviruses. In Numb KO cells, Hh signaling is turned on by cilium-PKI to the levels comparable to those in WT cells. Hh signaling activity is assessed by Gli1 transcript levels. This condition corresponds to the scenario **a_3_**. Statistics in (**b**, **c**): One-way ANOVA with multiple comparisons (Tukey test). (**d**): Two-way ANOVA with multiple comparisons (Tukey test). All error bars represent SD, n=3. ****, p < 0.0001; ***, p < 0.001. ns, not significant.

**Supplementary Figure 11.**
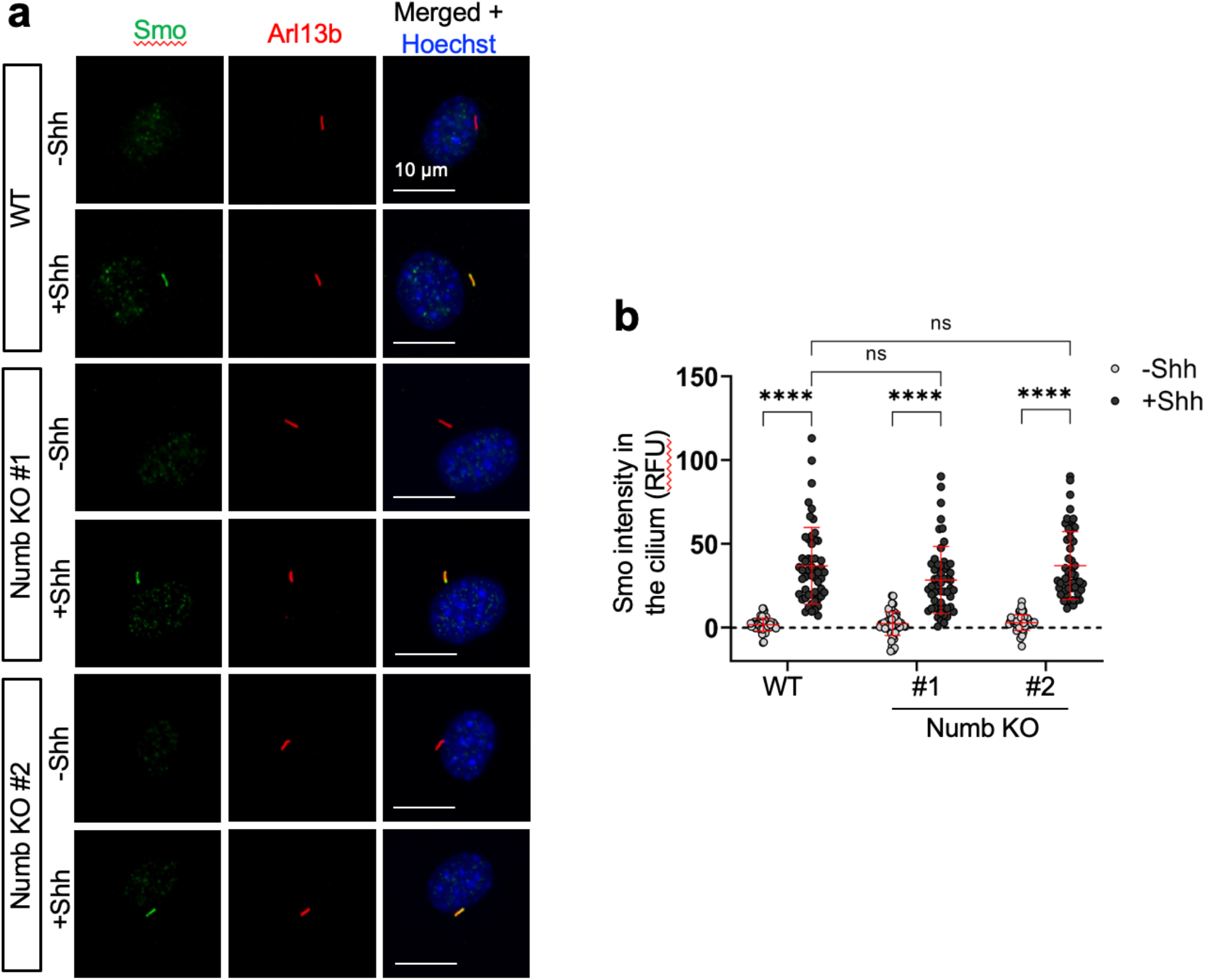
The effect of Numb loss on ciliary accumulation of Smo. (**a**) Immunofluorescence of Smo in the cilium in WT and Numb KO cells. Cells were serum-starved with or without Shh in the culture medium for 24 h before staining. (**b**) Quantification of Smo fluorescence intensity in the cilium. Data are shown as mean ± SD. n = 50 cilia were quantified per condition. RFU, relative fluorescence unit. Statistics: Two-way ANOVA with multiple comparisons (Tukey test), ****, p < 0.0001. ns, not significant.

**Supplementary Figure 12.**
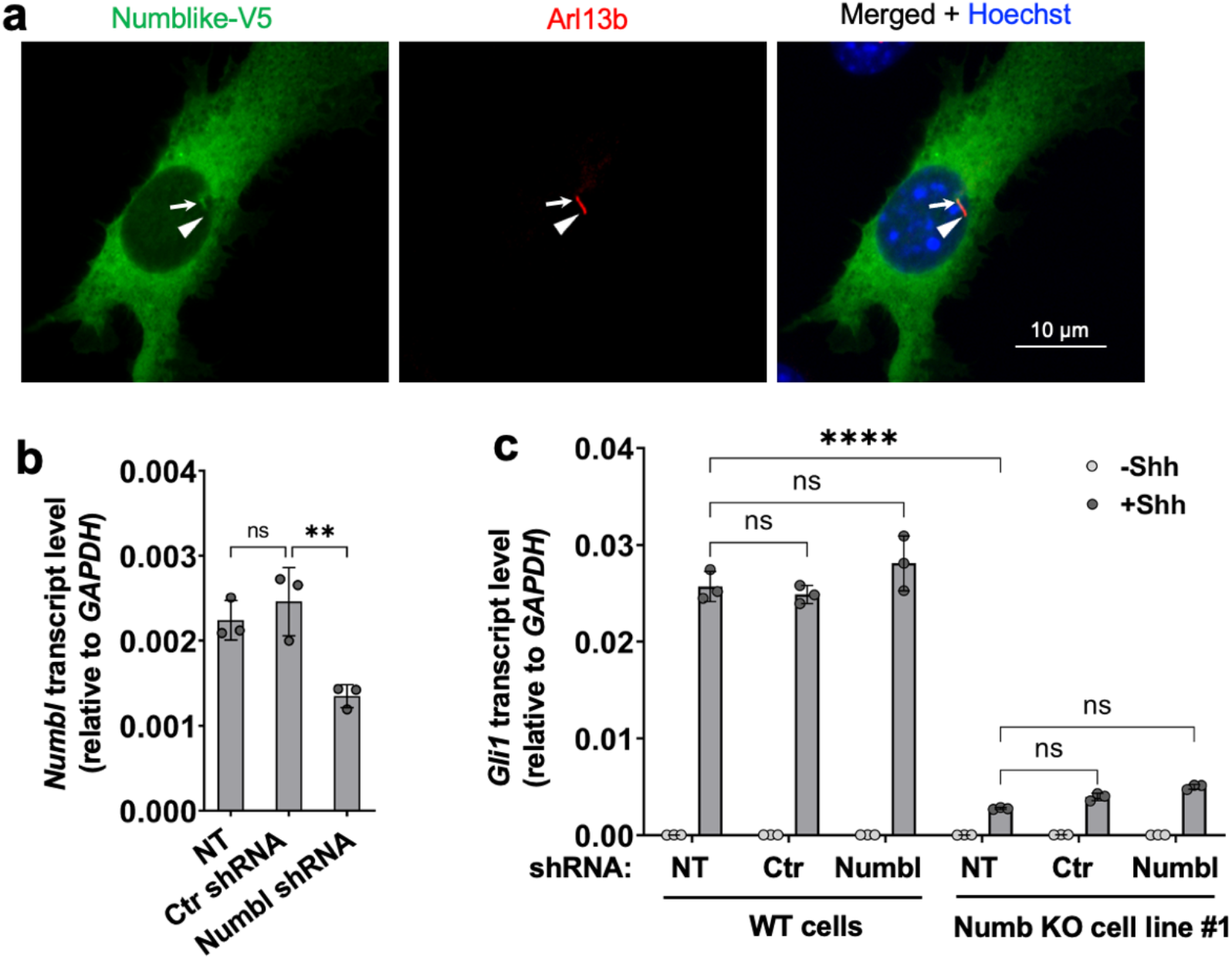
Numblike knockdown does not impact Hh signaling. (**a**) Numblike-V5 was expressed in NIH3T3 cells and co-immunostained with the cilium marker Arl13b. Note that in comparison to Numb (Fig 3a), Numblike diffuses uniformly throughout the cytosol, without displaying punctate localization. (**b**) Numblike shRNA significantly reduced Numbl gene expression in NIH3T3 cells. Numblike mRNA levels were measured by qPCR. (**c**) Parental (WT) NIH3T3 cells or Numb KO cells were infected with lentiviruses that express Numblike shRNA. 3 days after infection, cells were serum-starved for 24 h with or without Shh. Hh signaling activity was evaluated via qPCR measuring the transcript levels of Gli1. Control (Ctr) shRNA is a non-mammalian targeting shRNA. NT: non-transfected. Data are shown as mean ± SD, n=3. Statistics: one-way ANOVA in (**b**) and Two-way ANOVA with multiple comparisons (Tukey test) in (**c**). ****, p < 0.0001; **, p < 0.01; ns, not significant.

**Supplementary Figure 13.**
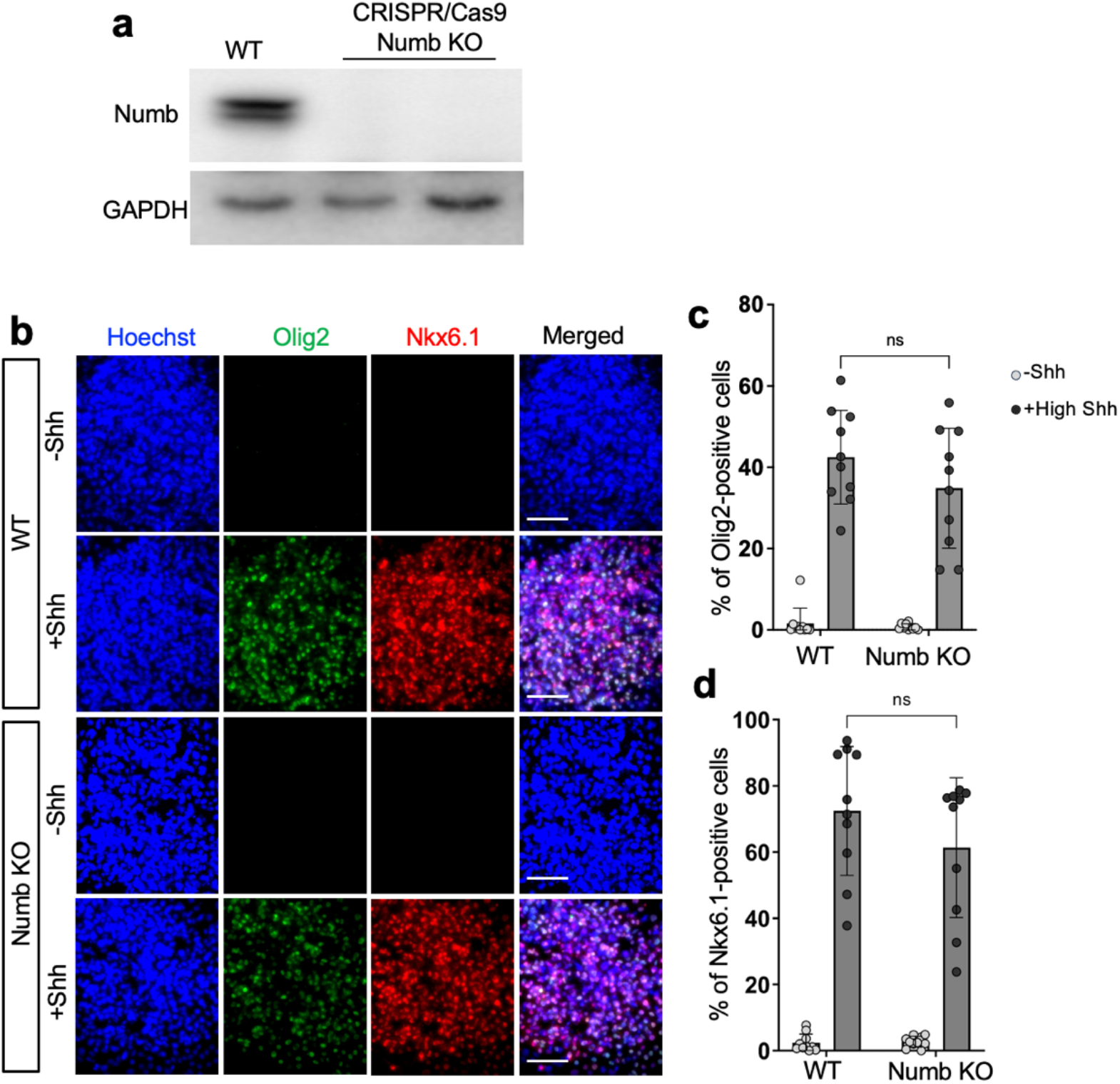
Numb loss has moderate or no impact on the differentiation of NPCs that is reliant on medium to low Hh signaling activity. (**a**) Western blotting results show that Numb protein was ablated in mouse embryonic stem cells. (**b-c**) Representative images of NPC differentiation into Nkx6.1- and Olig2-positive neural progenitors after 3 days induction. The total of 15 images from different colonies were quantified. Each image represents one NPC colony consisting of approximately 200–300 cells. Each dot represents the data from one image of an NPC colony. Statistics: Two-way ANOVA with multiple comparisons (Tukey test). All error bars represent SD. *, p < 0.05; ***, p < 0.001; ns, not significant.

**Supplementary Figure 14.**
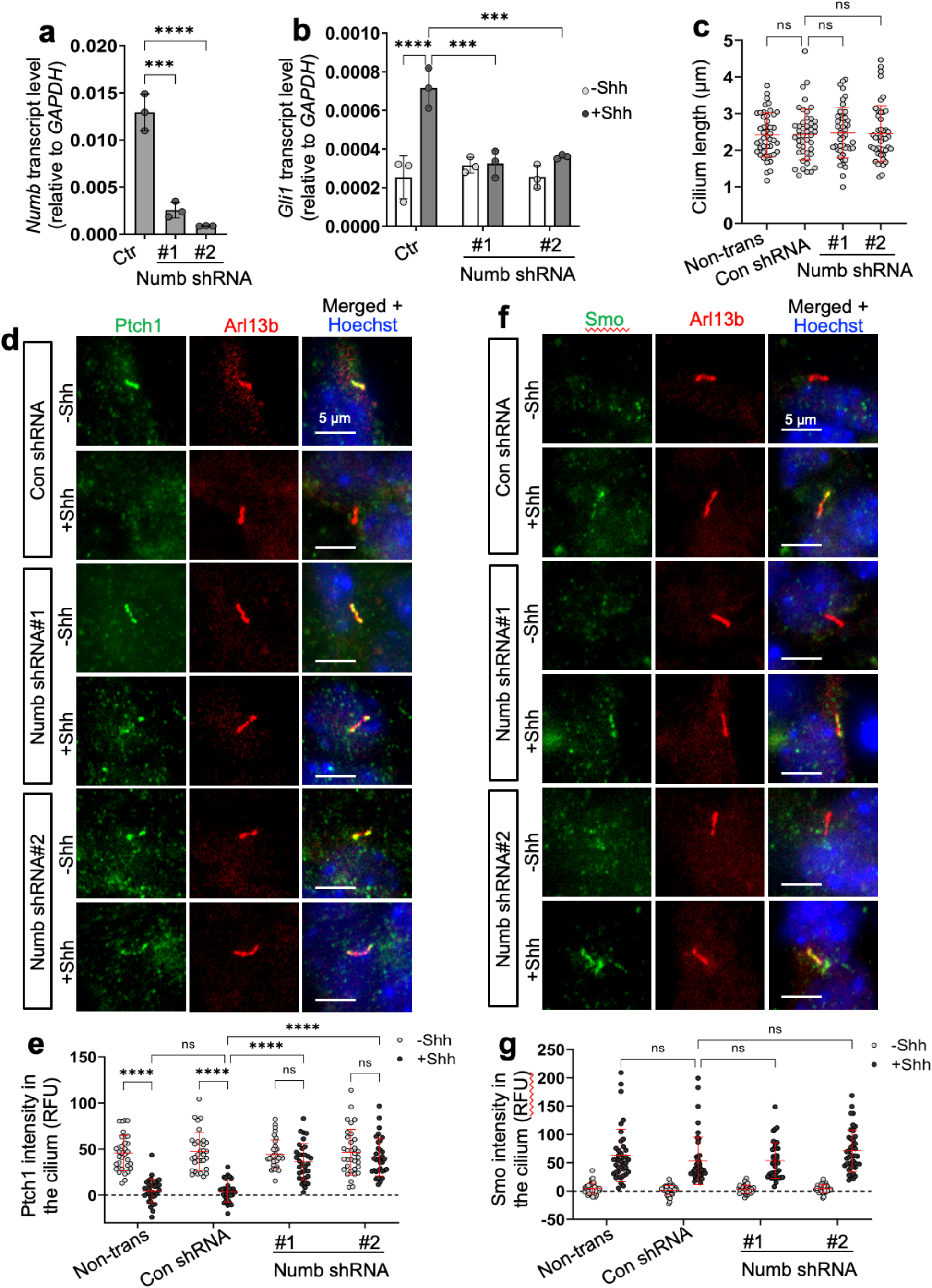
Numb is required for activation of Hh signaling and Shh-induced Ptch1 exit from the cilium in GCPs. (**a**) Numb shRNAs significantly reduced Numb gene expression in primary cultured GCPs. Numb mRNA levels were measured by qPCR. (**b**) Numb knockdown reduced Shh-induced Hh target gene transcription. 3 days after lentivirus-mediated transfection, GCPs were stimulated for 24 h with or without 1 μg/ml Shh. (**c**) Quantification of the cilium length in non-transfected, control shRNA or Numb shRNA transfected GCPs. 3 days after lentivirus-mediated transfection, GCPs were immuno-stained with Arl13b. n=45 cilia were quantified per condition. (**d**) Immunofluorescence staining of endogenous Ptch1 in the cilia of primary cultured GCPs. 3 days after lentivirus-mediated transfection, GCPs were pretreated with SAG for 24hr to induce Ptch1 expression. After that cells were stimulated with 1 μg/ml Shh or control medium for 1 h before being fixed for immunostaining. Scale bar, 5 µm. (**e**) Quantification of endogenous Ptch1 intensity in the cilia of primary cultured GCPs. n=30 cilia were quantified per condition. (**f**) Immunofluorescence of Smo in the cilia of primary cultured GCPs. 3 days after lentivirus-mediated transfection, GCPs were treated for 24 h with or without 1 μg/ml Shh. Scale bar, 5 µm. (**g**) Quantification of Smo fluorescence intensity in the cilium. n=40 cilia were quantified per condition. Statistics in (**a, c**): One-way ANOVA with multiple comparisons (Tukey test). (**b**, **e, g**): Two-way ANOVA with multiple comparisons (Tukey test). All error bars represent SD. ***, p < 0.001; ****, p < 0.0001. ns, not significant.

**Supplementary Figure 15.**
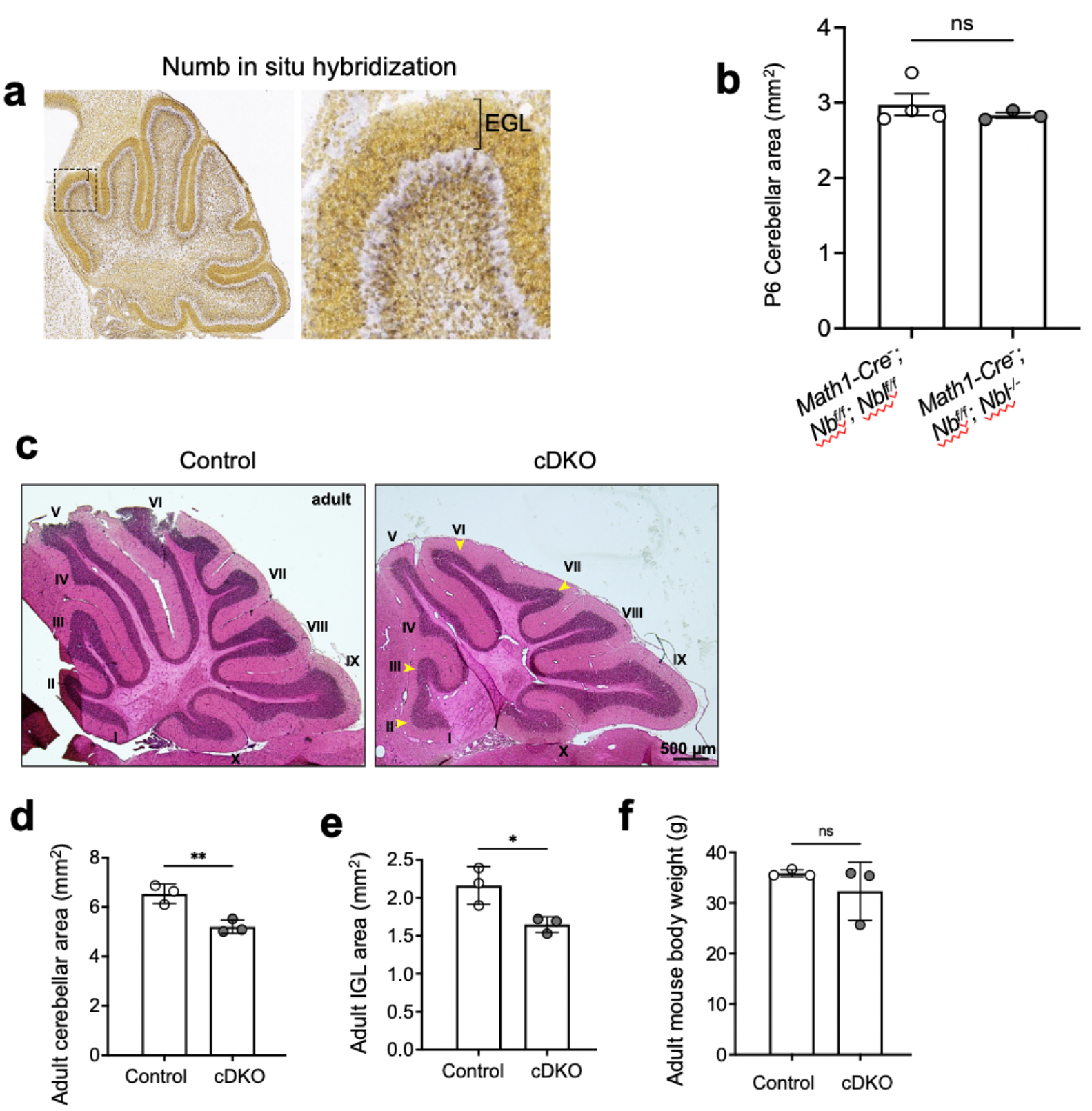
Numb/Numbl cDKO leads to reduced cerebellar size in adult mice. (**a**) In situ hybridization on sagittal mouse sections at P4 for Numb. Numb mRNA is present in the EGL, Purkinje cells, and IGL. Bracket denotes the EGL. Box indicates zoomed area. Images are obtained from The Allen Brain Atlas. (**b**) Quantification of the overall cerebellar area in P6 mice measured at the same mediolateral level. n = 3 cerebella per group. (**c**) Sagittal sections of cerebellum from adult control and *Numb;Numbl* cDKO mice with H & E staining. Roman numerals indicate the lobules and arrowheads indicate under-developed lobules in *Numb;Numbl* cDKO cerebella. (**d-e**) Quantification of the overall cerebellar area and IGL area in adult mice measured at the same mediolateral level. n = 3 cerebella per group. (**f**) Body weight in adult mice. n = 3 mice per group. All results are presented as mean ± SEM. Statistics: Student’s t-test, *, p<0.05; **, p<0.01. ns, not significant.

## ACKNOWLEDGEMENTS

We thank Dr. Rajat Rohatgi (Stanford University) for the gifts of CRISPR/Cas9 reagents, and Dr. Matthew Scott lab for the antibodies against Ptch1 and Smo. We thank Dr. David Gravano (Stem Cell Instrumentation Foundry, University of California, Merced) for fluorescence-activated cell sorting, and Dr. David Ardell (University of California, Merced) for consultation of proteomic data analysis. We thank Dr. Christopher J. Westlake (Center for Cancer Research, National Cancer Institute) for the gift of EHD1-GFP plasmid. We thank Francois Depault, Christine Jolicoeur and Jessica Barthe for expert technical assistance. We thank Julien Ferent for critical discussions and Kevin Zhang for help with the cerebellum histological analysis. The data in this work was collected, in part, with a confocal microscope acquired through the National Science Foundation MRI Award Number DMR-1625733. Mass spectrometry was performed in the Vincent Coates Foundation Mass Spectrometry Laboratory, Stanford University Mass Spectrometry (RRID:SCR_017801) by Dr. Ryan Leib, Dr. Fang Liu, Kratika Singhal, and Rowan Matney. Correlative ImmunoEM was performed at the Electron Microscope Lab at University of California, Berkeley, under the supervision of Danielle Jorgens. Research in the laboratory of X.G. was supported by NIH/NCI R15 (CA235749), NIH R01 (GM143276), NIH R21 (CA274595), and NSF CAREER award (IOS-2143711). Research in the laboratory of F.C. was supported by the Canadian Institutes of Health Research (FDN334023 and PJT471325), Brain Canada-Weston Foundation (to F.C. and M.C.), and the Canada Foundation for Innovation (33768). F.C. holds the Canada Research Chair in Developmental Neurobiology.

## AUTHOR CONTRIBUTIONS

X. Ge, X. Liu, P.T. Yam and F. Charron conceived the project. X. Ge and X. Liu designed the experiments on cilium proteomics and Numb mechanistic study in cultured cells. X. Liu performed the experiments, analyzed, and interpreted the data on cilium proteomics and the Numb characterization in cultured cells. E. Cai curated and characterized the Numb CRISPR/Cas9 knockout cell lines. O. T. Gutierrez and J. Zhang performed experiments related to quantifications in the cilium length and cilium protein levels in NIH3T3 cells and in primary cultured GCPs. J. H. Kong designed and supervised experiments in NPCs. R. Gen did the CRISPR/Cas9 Knockout in mESC cells and genotyped the colonies. T. Marks performed the NPC differentiation, performed qPCRs and the immunostaining staining and imaging in differentiated NPCs. P.T. Yam, W.J. Chen, S. Schlienger, V. Jimenez Amilburu, V. Ramamurthy, M. Cayouette and F. Charron designed and performed the experiments on numb;numbl knockout mice, and analyzed and interpreted the data on cerebellar development and GCP proliferation. T. Branon and A. Ting provided the reagents for TurboID and provided technical support for TurboID related proteomics. X. Ge, X. Liu, P. T. Yam and F. Charron wrote the paper.

## MATERIALS AND METHODS

### Cell line generation, cultivation, and manipulation

NIH3T3 cells were cultured in DMEM (supplemented with 10% calf serum). Ciliation was induced by reducing the growth media to 0.5% CS for 24 h. Transfections were performed using Lipofectamine 2000 (Invitrogen) according to manufacturer’s guidelines. Numb gene was disrupted in NIH3T3 cells using CRISPR/Cas9-mediated genome editing with gRNAs targeting exon 1. Clones of cell line were obtained by single-cell sorting. Clones with the disrupted gene were screened by immunofluorescence (IF) and Western blotting for the target protein using protein-specific antibodies. Selected positive clones were further characterized by sequencing, confirming missense mutation leading to early termination of translation and frameshift mutation. NIH3T3 cell lines stably expressing Cilium-TurboID and Non-cilium-TurboID were generated using lentivirus plasmids encoding Cilium-TurboID and Non-cilium-TurboID and confirmed by sequencing. To induce Shh signaling, growth media were supplemented with either 100 nM SAG or ShhN conditioned medium (20-30% [vol/vol] depending on batch) produced with 293 EcR-Shh-N cells.

### TurboID labeling experiments and subcellular fractionation

Biotin labeling experiments were performed as described^22^. Briefly, cells were incubated in the presence of 50 μM biotin for 10 min. For non-labelled samples, water was added instead of biotin. After 10 min of incubation at 37°C, the medium was aspirated quickly, and cells were washed three times with 1× PBS. For fluorescence microscopy, cells were immediately fixed. For proteomic and Western blot analyses, cells were lysed with subcellular fractionation buffer (20 mM HEPES, 10 mM KCI, 2 mM MgCl2, 1 mM EDTA, 1 mM EGTA, 1mM DTT, pH 7.5 and protease inhibitors) and by 15 passages through a 27-G needle. Nuclei were separated from the post-nuclear supernatant (PNS) by centrifugation at 720 x g (3,000 rpm) for 5 min The PNS was also respun at 8,000 rpm (10,000 x g) for 5 min and then brought to Lysis Buffer (0.5% NP-40, 0.1% SDS, 0.5% sodium deoxycholate, 150 mM NaCl, 25 mM Tris/HCl, pH 7.5, and protease inhibitors), sonicated and clarified by centrifugation at 10,000g for 10 min.

### Streptavidin capture

After determining the protein concentrations of lysates from the TurboID labeling experiments, they were adjusted to equal concentrations and volumes as starting material, from which samples were taken as loading control for SDS-PAGE and Western blot analysis. The remaining lysates were added onto washed and equilibrated streptavidin magnetic beads, and biotinylated proteins were allowed to bind for 1.5 h at room temperature. Beads with bound proteins were washed extensively with a series of buffers to remove nonspecific binders: twice with Lysis Buffer, once with 1 M KCl, once with 0.1 M Na2CO3, once with 2 M Urea in 10 mM Tris-HCl, pH 8.0, twice with Lysis Buffer, and finally with 1 x PBS. For Western blot analysis, the beads were eluted with 2 x SDS sample buffer.

### On-beads trypsin digestion of biotinylated proteins and TMT labeling

Beads with bound proteins were incubated in 100 mM TEAB and reduced using 10 mM DTT at 55°C for five minutes, followed by 25 minutes at room temperature with head-over-head mixing. The reduced proteins were alkylated using 30 mM acrylamide for an additional 30 minutes of room temperature with head-over-head mixing. Peptides were digested using trypsin/LysC (Promega) overnight with head-over-head mixing. Peptides were quenched with 1% formic acid and separated from the beads; 0.1% formic acid was used as a second elution prior to C18 microspin column clean up and speedvac to dryness. Peptides were quantified using the Pierce Quantitative Fluorometric Peptide Assay (Thermo) to match input peptide amounts for TMT labeling. Aliquots of each sample were pooled for a common reference prior to isobaric labeling. Peptides were resuspended in 100 mM TEAB and tagged as described in the manufacturer’s protocol. Tagged peptides were then pooled to generate a multiplexed sample. This sample was then fractionated offline using high pH reverse-phase fractionation to increase peptide observations.

### Mass spectrometry

In a typical mass spectrometry (MS) experiment, dried, desalted, labeled peptides were reconstituted in 2% aqueous acetonitrile. MS experiments were performed using liquid chromatography (LC) with an Acquity M-Class UPLC (Waters) followed by MS using an Orbitrap Fusion Tribrid MS (Thermo Scientific). A flow rate of 300 nL/min was used, where mobile phase A was 0.2% (v/v) aqueous formic acid and mobile phase B was 0.2% (v/v) formic acid in acetonitrile. Analytical columns were prepared in-house, with an internal diameter of 100 μm packed with NanoLCMS solutions 1.9 um C18 stationary phase to a length of approximately 25 cm. Peptides were directly injected into the analytical column using a gradient (2% to 45% B, followed by a high-B wash) of 120 min for each fraction. MS was operated in a data-dependent fashion using Collision-Induced Dissociation (CID) fragmentation for MS/MS spectra generation collected in the ion trap on the Fusion with MS3 detection of the reporter ions using synchronous precursor selection (SPS) to improve measuring accuracy as described previously^72^.

### MS Data processing

For data analysis, the .RAW data files were checked using Preview (Protein Metrics) to verify calibration and quality prior to further analysis. Data were then processed using Proteome Discoverer v1.5 (Thermo) for quantitative analysis of reporter ion ratios, and using the Byonic (Protein Metrics) node to identify peptides and infer proteins against the Mus musculus database from Uniprot (including isoforms and concatenated with common contaminant proteins). Proteolysis with Trypsin/LysC was assumed to be specific with up to two missed cleavage sites, and allowing for common modifications. Precursor mass accuracies were held within 12 ppm with fragment ions held within 0.4 Da (for CID) and 12 ppm (for HCD). Proteins were held to a false discovery rate of 1%, using standard approaches described previously (Elias and Gygi, 2007).

### Numb KO mESC generation and neural progenitor differentiation

The Numb gene was disrupted in a stable Gli-binding site (GBS)-Venus Hedgehog signaling reporter line (DV12) generated in HM1 mouse embryonic stem cells (mESCs)^52^. To ablate Numb function, CRISPR/Cas9-mediated genome editing was used. Briefly, two sequences targeting exon 1 of Numb (5’-GAAAGACGTTTATGTCCCAG-3’ and 5’-GGAAGCTACACTTTCCAGTG-3’) were individually cloned into plasmid pSpCas9(BB)-2A-Puro (PX459) V2.0 (Addgene #62988). These guide constructs were then electroporated into mESCs using the Lonza Nucleofector 2b Device and Cell Nucleofector Kit (Lonza #VAPH-1001). mESCs were cultured in feeder-free 2i media [DMEM/F-12 (Gibco #11320033) and Neurobasal Medium (Gibco #21103049) (prepared at a 1:1 ratio) supplemented with N-2 Supplement (Gibco #17502048), serum free B-27 Supplement (Gibco #17504044), penicillin/streptomycin (Gibco #15140122), glutaMAX (Gibco #35050061), bovine serum albumin (Thermo Scientific Chemicals #AAJ64248AE), 55 μM 2-mercaptoethanol (Gibco #21985023), 3 μM CHIR99021 (Axon #1386), 1 μM PD 98059 (Tocris Bioscience #1213), and ESGRO recombinant mouse LIF protein (1000 units/ml, MilliporeSigma #ESG1107)]. Antibiotic selection was initiated 24 hours after nucleofection in 2i media containing 1.5 μg/ml puromycin (Fisher BioReagents #BP2956100) for 72 hours. Clones were picked individually, expanded, and genomic DNA was collected using QuickExtract DNA extraction solution (Lucigen, #QE09050). The region surrounding the targeted site was PCR-amplified (F: 5’-agggtttggggtgggtttt-3’ and R: 5’-gctttgtctgggttccttcc-3’) to visualize the successful deletion facilitated by the guides. A loss of full-length Numb protein was then verified by Western blotting using protein-specific antibodies.

The mESCs were differentiated into spinal neural progenitors using a previously described protocol by Pusapati et al.^52^, which was modified with minor modifications from Gouti et al.^51^. To prepare the mESCs for differentiation, the mESCs were passaged twice on CF1 mitomycin C-treated mouse embryonic fibroblasts (MitC-MEFs, Gibco Cat #A34959) in mESC media [DMEM containing high glucose (Cytiva #SH30081.FS), 15% fetal bovine serum (Gibco #A5256701), MEM non-essential amino acids (Gibco #11140076), penicillin/streptomycin (Gibco #15140122), glutaMAX (Gibco #35050061), EmbryoMax nucleosides (MilliporeSigma #ES008D), 55 μM 2-mercaptoethanol (Gibco #21985023), and ESGRO recombinant mouse LIF protein (1000 units/ml, MilliporeSigma #ESG1107)]. For spinal neural progenitor differentiation, the feeders were first removed from the mESCs. Removal of MitC-MEFs was accomplished by first lifting all the cells off of the plate using 0.25% trypsin/EDTA (Gibson #25200072), inactivating the trypsin using mESC media (supplemented with no LIF), and then incubating the cells on 10 cm tissue culture plates for two short (20 min) successive periods. To induce differentiation, mESCs were plated onto either coverslips or CellBIND plates (Corning #3335) treated with 0.1% gelatin (Sigma Aldrich Fine Chemicals Biosciences #G139320ML) and cultured in N2B27 media [DMEM/F-12 (Gibco #11320033) and Neurobasal Medium (Gibco #21103049) (prepared at a 1:1 ratio) supplemented with N-2 Supplement (Gibco #17502048), serum free B-27 Supplement (Gibco #17504044), penicillin/streptomycin (Gibco #15140122), glutaMAX (Gibco #35050061), bovine serum albumin (Thermo Scientific Chemicals #AAJ64248AE), 55 μM 2-mercaptoethanol (Gibco #21985023), and varying components. On Day 0 (the day the cells were plated) and Day 1, the cells were cultured in N2B27 with 10 ng/ml mouse basic fibroblast growth factor (bFGF) recombinant proteins (R&D systems #313FB025/CF). On Day 2, the media was changed to N2B27 with 10 ng/ml bFGF and 5 μM CHIR99021 (Axon #1386). On Day 3, the cells were cultured in N2B27 with either no SHH [100 nM retinoic acid only (RA, Sigma Aldrich Fine Chemicals Biosciences #R262550MG)] or high SHH [100 nM RA and 100 nM SHH). No Shh and High Shh treatments were continued for 3 days before harvesting.

### Quantitative real-time PCR

NIH 3T3 cells were grown in 24-well plates in regular growth medium for 24 h, followed by serum starvation to induce ciliation and addition of SAG or Shh-N conditioned (or control) medium to induce Shh signaling. After 24 h, cells were RNA extracted using Trizol reagent (Invitrogen). Quantitative PCR was performed on 100 ng total RNA per reaction using Quantstudio 3 System (Applied Biosystems) and qPCR reagents (QuantaBio; qScript XLT One-Step RT-qPCR ToughMix, Low ROX). The TaqMan gene expression probes used were Mm00477927_m1 (Numb), Mm00477931_m1 (Numblike), Mm00489385_m1(Sufu), Mm00494645_m1 (Gli1), Mm00436026_m1 (Ptch1) and Mm99999915_g1 and GAPDH to normalize the samples.

For the analysis of the neural progenitor cells (NPCs), RNA extraction, complementary DNA synthesis, and qRT-PCR analysis were conducted as previously described^51, 52^. The following primers were used for qRT-PCR: Gli1 (F: 5’-ccaagccaactttatgtcaggg-3’ and 5’-agcccgcttctttgttaatttga-3’), Nkx6.1 (F: 5’-cccggagtgatgcagagt-3’ and R: 5’-gaacgtgggtctggtgtgtt-3’), Olig2 (F: 5’-agaccgagccaacaccag-3’ and R: 5’-aagctctcgaatgatccttcttt-3’), Nkx2.2 (F: 5’-cagcctcatccgtctcac-3’ R: 5’-tcacctccatacctttctcc-3’), Ptch1 (F: 5’-tgacaaagccgactacatgc-3’ and R: 5’-agcgtactcgatgggctct -3’). Transcript levels were calculated relative to Gapdh (F: 5’-agtggcaaagtggagatt-3’ and R: 5’-gtggagtcatactggaaca-3’) using the ΔΔCt method.

P6 GCPs were seeded at 2.5×10^6^ cells/well in 6-well plates. GCP cells were treated or not with 1 nM recombinant ShhN for 20 h and pellets were directly resuspended in the lysis buffer. RNA was purified using the RNeasy Mini Plus kit (QIAGEN). cDNA was synthesized using the Transcriptor First Strand cDNA synthesis kit (Roche) using total RNA. Real-time PCR mixes were prepared using Perfecta SYBR Green Supermix (Quanta Biosciences). Reactions were performed in triplicate, and the amount of cDNA per reaction was 5 ng. Results were analyzed using the Comparative Ct method. The levels of Gli1 mRNA were normalized to Gusb mRNA. The primers used were Gli1 F: GCA GTG GGT AAC ATG AGT GTC T and Gli1 R: AGG CAC TAG AGT TGA GGA ATT GT.

### Immuno-electron microscopy

For correlative light and electron microscopy (CLEM), NIH3T3 cells expressing Numb-HA were plated on gridded-glass bottom dishes (MatTek, cat. no. P35G-1.5-14-C-GRD). After 4% PFA fixation, cells were permeabilized with 0.2% Triton X-100 in PBS for 10min and underwent immunostaining with Rabbit anti-HA (cell signaling, Cat # 3724), follow by the 2^nd^ antibody of Alexa 488 FluoroNanogold-conjugated anti-rabbit IgG (Nanoprobes, #7204). The nanogold signal was enhanced via the GoldEnhance EM Plus kit (Nanoprobes, #2114). After that, fluorescence and DIC images were taken with a Leica DMi8 microscope. The position of cells was recorded using grid numbers on cover glasses. We used a modified protocol from UC Berkeley EM facility to process the cells. Briefly, cells were post-fixed in 1% OsO_4_ with 1.6% potassium ferricyanide (K_3_Fe(CN)_6_) in PBS for 30 minutes. Cells were then rinsed with 1xPBS 3 times for 5 minutes each and briefly rinsed with distilled H_2_O. To dehydrate the cells, an increasing percentage of 200 proof ethanol was added to the plates in 10-minute increments: 30%, 50%, 70%, 95%, 100%, 100%, 100%. Next, cells were infiltrated with increasing resin amounts to ethanol for 30 minutes each; 1:4, 1:2, 1:1, 2:1. Finally, cells were incubated with pure resin 3 times for 30 minutes each. At the end of incubation, the remaining resin was removed, and a thin layer of pure resin was added to cover the well containing cells. The samples were then incubated at 60°C for 16 hrs and microtomed for imaging.

### Immunofluorescence and microscopy

Cells were grown on round 12 mm #1.5 coverslips and fixed in 4% PFA for 10–15 min at room temperature. After fixation, cells were permeabilized and blocked in blocking buffer (2% donkey serum, 0.2% triton X-100, in PBS) for 1 h at room temperature. After blocking, cells were incubated with primary antibody diluted in blocking buffer for 1 h at room temperature or 4°C overnight, washed three times with 1× PBS over 15 min, and incubated with Alexa Fluor 488-, rhodamine-, or Alexa Fluor 647-coupled secondary antibodies (Jackson ImmunoResearch) in blocking buffer for 1 h, and then incubated in Hoechst (1ug/ml in PBS) for 10 min. Finally, cells were washed five times in PBS and mounted on glass slides using Fluoromount G (Southern Biotech; 0100-01). Prepared specimens were imaged on a Leica DMi8 (LAS X software) with Plan Apochromat oil objectives (63x, 1.4 NA) or a LSM880 confocal microscope (Zeiss). Images were processed using ImageJ.

### Expansion microscopy

The gelation and proteinase digestion steps are similar to the proExM protocol by Tillberg et al.^40^, with two modifications: (1) after cells are fixed and stained with primary antibodies, we added 4%PFA to cross link the antibody to the samples; and (2) we stained the sample with secondary antibody after proteinase digestion, with the intention to protect the fluorescent dyes.

Briefly, fixed cells were incubated in primary antibodies for 2 h at room temperature or at 4°C, overnight and then were fixed in 4% PFA for 20 min. After fixation, cells were incubated in the 0.1 mg/ml AcX (Acryloyl-X SE, Invitrogen, cat. no. A20770) solution for 2 to 3 hr at room temperature. Then cells were polymerized in the gelling solution by mixing Stock X (8.6% Sodium acrylate, 2.5% Acrylamide, 0.15% N,N’-Methylenebisacrylamide, 11.7% Sodium chloride, 1x PBS), water, 10% TEMED stock solution, and 10% APS stock solution in a 47:1:1:1 ratio. The gel was then digested with proteinase K (NEB, cat. no. P8107S) at a final concentration of 8 U/ml in digestion buffer (50 mM Tris, pH 8.0, 1 mM EDTA, 0.5% Triton X-100, and 0.8 M NaCl) for 12 h at room temperature. After digestion, proteinase K was removed by four washes with excessive PBS (10 min each time) and the gel was incubated in the secondary antibody for 2 h at room temperature. The post-expansion labeled hydrogel was then washed and expanded by four washes with excessive water (at least 30 min each time) and mounted with superglue. Imaging was performed by a AiryScan LSM880 confocal microscope (Zeiss) with a 63x lens.

During imaging, the elastic nature of the hydrogel makes it difficult to complete a z-stack optical scanning. The short working distance of the lens presses the hydrogel, dislocates the samples’ position and causes image blurring. To avoid the potential artifacts of imaging blur, we captured one single optical plane focusing only on the ciliary pocket.

### Quantification of cilium length and protein intensity in the cilium

Cilium length was measured in Leica LAS X software. A line was drawn along the fluorescent signal corresponding to the ciliary marker, and the length of this line was defined as the length of the primary cilium. The ciliary protein intensity was measured in ImageJ. Briefly, we first outlined the contour of an individual cilium in the channel of cilium staining. After that, the fluorescence intensity within this contour was read out for each individual channel of the corresponding protein (L1). Finally, the individual cilium contour was manually dragged to the region right next to the cilium, and the intensity within the contour was read out as the background (L2). The final fluorescence intensity for that channel was defined as L= L1 - L2.

### Quantification of Ptch1 incorporation in Numb-containing CCVs

The quantification was done in Fiji. All quantified images are done on confocal microscope images. The region of interest (ROI) was defined as a circle with the cilium base as the center and the cilium length as the radius. Within the ROI, we remove the background by subtracting the global threshold value. Thresholded Manders’ Coefficients of Ptch1-YFP was calculated using BIOP JACoP plug-in in ImageJ, based on the co-localization methods described in Manders et al^73^.

### SDS-PAGE and Western blotting

Standard techniques were used for SDS-PAGE and Western blotting. Cells were first washed with PBS, and then were scraped off from the culture surface in RIPA buffer (1% NP-40, 0.1% SDS, 0.5% sodium deoxycholate, 150 mM NaCl, 25 mM Tris/HCl, pH 7.5, and protease inhibitors). Lysates were cleared by centrifugation (10,000 ×g at 4°C for 15 min), and 25 µg protein was separated on 10% SDS-PAGE gels and transferred onto PVDF membranes. After blocking in 5% BSA, membranes were washed and incubated with primary antibodies and secondary antibodies. Finally, proteins were detected with chemiluminescence substrates (Thermo Fisher Scientific; 34076). Quantitation of bands was performed using ImageJ.

### Animals

All animal work was performed in accordance with the Canadian Council on Animal Care Guidelines and approved by the IRCM Animal Care Committee. Mice were maintained in the IRCM specific pathogen-free animal facility. All mouse lines have been previously described: Math1-Cre^74^, numb conditional allele and numbl conditional allele^75^ (obtained from The Jackson Laboratory), and numbl null allele^76^. Mice of both sexes (not determined) were randomly used for experiments.

For experiments with conditional deletion of Numb and Numbl, control mice were *Math1-Cre^-^;Nb^f/f^;NbL^f/f^*, *Math1-Cre^-^;Nb^f/f^;NbL^f/-^*, or *Math1-Cre^-^;Nb^f/-^;NbL^f/-^*. Numb;Numbl cDKO mice were *Math1-Cre^+^;Nb^f/f^;Nbl^f/f^*.

### Histology

Brains were dissected and fixed by immersion overnight in 4% paraformaldehyde (PFA) diluted in PBS. The tissues were then cryoprotected in 30% sucrose, embedded in a sucrose:OCT (1:1) mix and frozen. Histology and immunochemistry were performed on sections as described previously^77, 78^. Rabbit anti-Numb antibody (Abcam 14140) was used at 1:250.

### Isolation of GCPs and in vitro proliferation assays

GCPs were isolated from P6 mice cerebella as described previously^57^. Briefly, isolated cerebella were cut into small pieces and treated with trypsin and DNase I. After trituration, single cell suspensions were centrifuged through a 30% to 65% percoll step gradient. Cells were harvested at the 30% interphase and then resuspended in Neurobasal supplemented with B27, 0.5 mM L-glutamine and penicillin/streptomycin and plated on plates precoated with 100 μg/ml poly-D-Lysine.

For the ^3^H-thymidine incorporation assay, GCPs were plated at 3 × 10^5^ cells per well in 96 well plates in triplicate and treated with ShhN for 48 h. GCPs were pulsed with 1 μCi/ml ^3^H-thymidine 12 hours after seeding the cells. The amount of ^3^H-thymidine incorporation was measured using the Filtermate harvester (PerkinElmer).

### Statistical analyses

Statistical analyses were performed with GraphPad Prism 8. For volcano plot graphs, Student’s t-test were used, and statistical analyses were performed in Prism 8. Hierarchical cluster analyses were performed according to Ward’s minimum variance method.

